# Integrating Semantic Retrieval, LLM-based Refinement, and Structured Expert Curation for Scalable AOP Gene Mapping

**DOI:** 10.64898/2026.06.25.734475

**Authors:** Alexandra Schaffert, Michele Fratello, Kaisla Kangas, Marcella Torres Maia, Giusy del Giudice, Lena Möbus, Cristina Accardi, Zeyad Al-Abdulraheem, Lorenzo Campini, Francesco Galardo, Antonio Federico, Giorgia Ciancaleoni, Hanna-Kaarina Juppi, Martin Paparella, Angela Serra, Dario Greco

**Affiliations:** Finnish Hub for Development and Validation of Integrated Approaches (FHAIVE), Faculty of Medicine and Health Technology, Tampere University, Tampere, Finland; Tampere Institute for Advanced Study, Tampere University,, Finland; University of Helsinki, Division of Pharmaceutical Biosciences, Faculty of Pharmacy, Helsinki, Finland; Institute for Medical Biochemistry, Medical University Innsbruck, Innsbruck, Austria

**Keywords:** Adverse Outcome Pathway, annotation, semantic similarity, large language model, toxicogenomics

## Abstract

Toxicogenomics can support regulatory toxicology, but its use is limited by the difficulty of translating molecular responses into mechanistic, decision-relevant interpretations. Adverse Outcome Pathways (AOPs) provide a framework for this translation, yet omics applications require scalable mapping of Key Events (KEs) to molecular features. Here, we present an AI-assisted, multi-step workflow for KE-to-gene mapping that uses embedding-based semantic retrieval to identify candidate ontology/pathway terms, large language model-assisted refinement to filter these candidates, and double-independent expert group curation with rule-based consolidation to finalize mappings and derive confidence scores. Compared with earlier NLP-based approaches, the workflow improves KE-to-ontology/pathway mapping performance and generates candidate annotations that better align with expert judgment while substantially reducing the need for manual augmentation. Explicit gene and protein mentions in KE titles were additionally grounded to improve specificity, and each curated mapping was assigned curator reason codes to support transparent, traceable, and confidence-aware reuse. Applied across AOP-Wiki, the workflow produced a comprehensive KE-to-gene set resource covering 1,254 KEs across 523 AOPs and linking 15,833 human genes. Utility is demonstrated through CTD-based AOP fingerprinting of curated reference chemical groups, highlighting expanded coverage and confidence-informed interpretation of chemical-associated gene signatures in an AOP context. The workflow and resulting resource provide a practical bridge between toxicogenomics and AOP-based mechanistic interpretation and support routine updating and future extension to additional omics layers within OECD Omics2AOP.

## 1. Introduction

Next-generation toxicology increasingly relies on high-dimensional molecular measurements, particularly transcriptomics and other omics readouts, to characterize biological perturbations induced by chemical exposures (Brockmeier et al., 2017; OECD, 2023b, 2025). However, despite clear potential, the use of toxicogenomics-based evidence in regulatory contexts remains constrained by the difficulty of translating complex patterns of molecular responses into mechanistically transparent and decision-relevant interpretations (Meier et al., 2024).

AOPs provide a suitable scaffold for this translation, linking an initial molecular perturbation through a series of measurable Key Events (KEs) to an Adverse Outcome (AO) relevant for human health or the environment (Ankley & Edwards, 2018). However, to integrate AOPs and omics data, the biological events described as KEs need to be linked to molecular features that can be monitored in omics datasets (Bakker et al., 2023; Brockmeier et al., 2017; Perkins et al., 2022; Saarimaki et al., 2023a). Gene sets derived from curated pathways, processes, and phenotypes offer a practical and widely adopted solution, enabling approaches that integrate experimental data with AOPs (Saarimaki et al., 2023b). In previous work, we established a curated link between AOP-Wiki KEs and gene-level representations (Saarimaki et al., 2023a) and showed that this enables practical AOP-informed analysis of molecular datasets, including AOP-based mechanistic interpretation, chemical grouping, comparison of biological experimental systems and species, and KE-anchored BMDs (Del Giudice et al., 2024; Maia et al., 2025; Saarimaki et al., 2023b; Serra et al., 2025).

Since that annotation effort, AOP-Wiki has expanded substantially, with many newly added AOPs and KEs. To remain useful as a shared framework for evidence integration, hypothesis generation, and mechanistic interpretation, KE-to-gene mappings require systematic updating in a way that is operationally scalable and supports routine expansion as the knowledge base grows (Bajard et al., 2023; Carusi et al., 2021).

The previous annotation approach (Saarimaki et al., 2023a) relied on traditional Natural Language Processing (NLP)-driven matching that prioritizes surface-level similarity (e.g., token overlap, keyword proximity, or other lexical similarity signals), which is typically effective when concepts are expressed using consistent terminology and relatively fixed phrasing. However, KE titles in AOP-Wiki frequently contain linguistically complex, highly contextual expressions and vary widely in phrasing and granularity (e.g., compositional mechanisms, implicit causal relationships, multi-clause descriptions, and domain-specific phrasing). Such complexity is known to challenge classical NLP pipelines (Turchin et al., 2023). As a result, lexically driven matching can yield many plausible but incorrect candidates (reduced precision) while missing valid matches expressed using different wording (reduced recall), increasing the need for time-intensive, manual expert review, which is an obstacle to scaling and broad adoption.

Recent advances in transformer-based language modeling offer a path forward. These models generate context-dependent representations of text, meaning that the interpretation of a word depends on the surrounding words rather than being fixed in all settings. As a result, models such as Bidirectional Encoder Representations from Transformers (BERT) can capture semantic similarity beyond direct word overlap (Gardazi et al., 2025). Large language models (LLMs) extend these capabilities. They are large Transformer-based generative models trained on massive corpora with an autoregressive “next-token prediction” objective, enabling them to follow natural-language instructions and perform new tasks via prompting rather than task-specific model redesign (Brown et al., 2020; OpenAI, 2023). This capability is particularly relevant for AOP annotation, where the core challenge is not only to retrieve semantically related terms, but determining whether a candidate pathway or ontology term is mechanistically consistent with a KE statement, including qualifiers such as directionality and biological context.

At the same time, ambiguity cannot be fully eliminated: KE phrasing may be underspecified, ontology coverage is incomplete, and multiple interpretations may be defensible. Because these mappings are reused across downstream applications, uncertainty should be made explicit rather than implicitly treating all KE annotations as equally reliable.

These considerations highlight the need for an annotation strategy that is scalable, meaning-aware, and transparent about uncertainty, aligning with broader efforts to improve machine-actionability and FAIR reuse of AOP-derived resources (Mortensen et al., 2025).

To address these needs, we developed a multi-step workflow that couples semantic candidate retrieval with expert-driven refinement and structured evaluation. Using this approach, we generated an updated, AOP-wide KE-to-gene annotation, with expanded coverage and an explicit confidence layer. These resources improve the scalability, consistency, and transparency of KE annotation and support reuse across downstream toxicogenomic applications.

## 2. Methods

### 2.1 Retrieval of KEs from AOP-Wiki for annotation

An XML snapshot of AOP-Wiki was downloaded (XML version 2025-04-01) and parsed to extract AOP, KE, and AO records, including stable identifiers and titles. KEs constituted the primary annotation unit. Entries with empty titles were excluded from automated annotation. The initial annotation scope comprised 1,503 KEs spanning all 530 AOPs present in AOP-Wiki at that time point. KE-level annotation was performed on this April 2025 snapshot because generation and expert curation of KE-to-gene mappings constituted the time-intensive core task of the workflow. To ensure that the final resource reflects a more current AOP-Wiki structure, KE presence and KE-to-AOP membership were subsequently updated using the AOP-Wiki XML snapshot dated 2026-04-01. KEs no longer present in the newer snapshot were removed from the final released AOP-level resource, and updated KE-to-AOP memberships were incorporated where already annotated KEs had become linked to additional AOPs. Because KE-level curation was attached to stable KE identifiers, this structural update did not require repetition of manual annotation.

### 2.2 Knowledgebase resources and term selection

Ontology and pathway term names and identifiers for GO, Reactome, WikiPathways, KEGG, and HPO were taken from MSigDB v2024.1.Hs to define a harmonized candidate term selection for semantic matching to AOP-Wiki KE titles. MSigDB provides curated, standardized collections of pathway/ontology terms assembled from multiple sources and delivered in a consistent naming/identifier framework, which we used to reduce downstream cleaning, and format harmonization during candidate generation. MSigDB v2024.1.Hs incorporates the following source snapshots for the relevant collections: GO (C5:GO, go-basic.obo 2024-04-24), Reactome (C2:CP:Reactome, v89), WikiPathways (C2:CP:WikiPathways, 2024-07-10), Human Phenotype Ontology (C5:HPO, 2024-04-26), and KEGG (C2:CP:KEGG; legacy pathway set). Importantly, MSigDB was used only to obtain standardized term labels/identifiers for candidate generation. After automated matching and subsequent group-curation finalized the accepted KE-term mappings, gene members were retrieved directly from the primary pathway/ontology sources for each accepted identifier, ensuring that gene sets reflect the most current content of the underlying resources at the time of retrieval rather than the MSigDB distribution (section 2.6).

### 2.3 Semantic similarity candidate retrieval

Candidate ontology/pathway terms for each KE were retrieved by assessing semantic similarity in embedding space between KE titles and knowledgebase term labels. Both were embedded using the pre-trained Sentence-BERT (SBERT) (Reimers & Gurevych, 2019) model all-mpnet-base-v2 . This model is based on the MPNet architecture (Song et al., 2020) and further optimized for semantic similarity tasks using a large and diverse corpus of more than one billion sentence pairs. This training objective is well aligned with the present task, which requires identifying semantically equivalent or closely related biological concepts between short phrases from different resources, namely KE titles and different ontology/pathway labels. Although biomedical domain specificity can be relevant, label-to-label entity matching depends particularly on robust semantic alignment in embedding space, including the ability to capture paraphrase-level similarity and variation in phrasing or abstraction. The chosen model is well suited to this setting because it has been extensively fine-tuned for sentence-level similarity across diverse text sources, including scientific abstracts and titles, supporting stable alignment between biologically meaningful phrases even when wording differs. Cosine similarity was computed between each KE embedding and all term embeddings, yielding a ranked candidate list per KE. Candidate terms with cosine similarity <0.6 were discarded, and the top 10 remaining candidates across all resources were passed to the subsequent refinement step. Similarity threshold choices were evaluated during technical validation (section 2.5).

### 2.4 LLM-based refinement

To further refine biologically meaningful ontology matches beyond similarity-based ranking, filtered candidate ontology sets were evaluated using a large language model (ChatGPT; GPT-4o, accessed programmatically via the OpenAI application programming interface (API) (OpenAI et al., 2024). For each KE, the API was queried with a structured prompt containing the KE identifier, KE name, and a comma-separated list of up to 10 candidate ontology/pathway terms. The model was instructed to act as a biomedical expert and to evaluate the biological and mechanistic relevance of each candidate term to the KE, explicitly considering both mechanistic fit and text similarity (Zenodo). The output format was constrained to return only the selected ontology terms as a comma-separated list, with a maximum of five retained terms; if no candidate was deemed biologically appropriate, the model was instructed to return “None”. To improve consistency of prompt interpretation across heterogeneous KE types, the prompt included seven few-shot examples. These comprised six positive/mixed examples, in which one or more ontology terms were retained from a list containing both relevant and irrelevant candidates, and one negative example in which no valid ontology term was available and the correct output was “None”. The examples were selected to cover a range of common annotation scenarios, including broad process-level KEs (e.g., inflammation, adipogenesis), receptor-/target-centric KEs (e.g., estrogen receptor agonism), phenotype-level KEs (e.g., ovulation, Leydig cell dysgenesis), sparse candidate lists, and cases in which only a single candidate should be retained despite the presence of semantically related distractors. In addition, the prompt explicitly instructed the model not to assume that earlier candidates in the list were more relevant than later ones, minimizing positional bias introduced by the SBERT ranking. Inference was performed with temperature = 0.35 and top_p = 1. In this setting, the low temperature reduces stochastic variation in output selection, whereas top_p = 1 disables nucleus truncation, such that all tokens remain available and sampling is not additionally constrained by probability-mass cutoffs. Together, these settings were chosen to favor stable and reproducible ontology selection over more variable or exploratory responses. The API output consisted only of the selected ontology identifiers, without explanatory text, and these were used as the final LLM-refined ontology annotations.

### 2.5 Technical validation during pipeline development

During workflow development, we used a random subset of 50 KEs to optimize the multi-step annotation pipeline and to obtain an initial technical assessment of retrieval and refinement performance. For the SBERT candidate retrieval step, we compared alternative candidate-selection strategies, including different similarity thresholds and top ranked candidates passed downstream. Retrieved candidates were manually labeled as mechanistically appropriate or inappropriate. Precision was defined as TP/(TP+FP), where TP denotes a relevant retrieved candidate and FP an irrelevant retrieved candidate. Recall was defined relative to a KE-specific manually curated reference set of relevant terms, TP/(TP+FN), where FN denotes a relevant term not retrieved under the evaluated configuration. Candidates below the similarity threshold were treated as not retrieved in the threshold-specific analysis. Based on this assessment, the final retrieval configuration was set to cosine similarity ≥ 0.6 combined with the Top 10 ranked candidates. The same 50-KE subset was also used to refine the LLM prompt and to obtain a technical assessment of the refinement step using the same manually curated relevance labels. Refinement quality was evaluated with rank-based metrics: Hit@1 (fraction of KEs for which the top-ranked selected term was relevant), Hit@5 (fraction of KEs for which at least one relevant term appeared among the top five selected terms), dynamic Hit@k (Hit@k with k matched to the number of relevant annotations per KE, bounded between 1 and 5), and MAP@k (mean average precision up to rank k, rewarding relevant selections appearing early). Because this subset was used to optimize both retrieval settings and prompting strategy, it should be regarded as a development subset rather than an independent final validation set.

### 2.6 Construction of KE gene sets

For each KE, the refined set of ontology/pathway terms was converted into a KE gene set by aggregating genes across the selected terms. Specifically, gene sets associated with each selected term were retrieved from the precompiled knowledgebase term-to-gene tables and mapped to Ensembl gene IDs; the KE gene set was defined as the union of these genes across all selected terms and resources. Provenance was retained by storing (i) the resource(s) contributing each term, (ii) the selected term identifiers per KE, and (iii) per-term gene set sizes, enabling resource-level decomposition of each KE gene set. Genes for each curated term ID were retrieved directly from the primary pathway/ontology sources and harmonized to human Ensembl gene IDs and gene symbols using Bioconductor org.Hs.eg.db v3.20.0 with AnnotationDbi v1.68.0 (Bioconductor 3.20; R 4.4.3), based on Ensembl and NCBI Entrez Gene annotation sources dated release 112 and 2024-09-20, respectively. GO term gene sets were obtained from the GOA human Gene Association File (GAF; GO release 2026-03-25) with negated (“NOT”) annotations excluded, and expanded using the GO go-basic.obo hierarchy (“go-basic” ontology snapshot used for propagation) via safe relations (is_a/part_of) to include genes annotated to descendant terms. HPO term gene sets were derived from phenotype_to_genes.txt (HPO release v2026-02-16), which provides phenotype-to-gene links that already include ancestor classes (supporting parent terms as collections of child terms). KEGG, Reactome, and WikiPathways gene sets were retrieved from their programmatic/downloadable resources (KEGG REST API, accessed April 2026; Reactome v96 released 2026-03-30; WikiPathways monthly GMT release 20260310) and merged into a unified term-to-gene mapping for downstream analysis. Across all annotated entities, the resulting compendium represented 15832 unique Ensembl genes.

### 2.7 Expert group curation and annotation scoring

To provide a scalable human quality-control layer, a structured expert group curation procedure was applied to KE–term mappings refined by the LLM step. In addition, mappings from Saarimaki et al. (2023a) were included in the same group-curation workflow and re-curated alongside the new candidate annotations. Prior to group curation, Saarimäki-derived KE-term mappings were filtered to remove obsolete terms and to exclude KEs that were no longer present in the more recent AOP-Wiki release. KEs with their proposed ontology/pathway annotations (including both workflow-derived candidates and Saarimäki et al. annotations) were randomly assigned to 11 curators, with each KE evaluated independently by two curators. Each KE–term mapping was assessed using a predefined three-level scheme: fit (term clearly represents the KE), partial (term captures the KE only partly or lacks sufficient specificity), or not_fit (term does not represent the KE). For partial and not_fit, curators selected one of 13 predefined reason codes (Supplementary file 1, Table S1). Curators additionally provided a binary inclusion decision indicating whether the mapping should be accepted (*yes*) or rejected (*no*) for KE gene set construction. Final decisions were consolidated using a rule-based procedure. If both curators agreed on the inclusion decision, that decision was adopted. In case of disagreement, outcomes were resolved based on the qualitative labels (fit vs partial was accepted; partial vs not_fit was rejected), while partial vs partial, fit vs not_fit, and other unclear combinations were referred for adjudication by a master curator. To support confidence-aware downstream use, a workflow-derived KE annotation quality score was calculated from curator labels across all finally accepted KE-term mappings for each KE, treating each curator label as an independent vote:

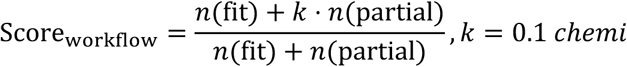

Scores ranged from 0.1 to 1.0 when at least one accepted mapping was present, with higher values indicating stronger curator agreement and higher annotation quality. The consolidated decision and score were stored alongside each KE-term mapping. In addition to workflow-derived annotations, curated single-gene associations derived from explicit gene/protein mentions in KE titles were incorporated into the final KE gene sets and treated as high-confidence evidence. To obtain a unified KE-level confidence measure, genes were assigned confidence values of 1.0 if supported by curated single-gene sources, or Score_workflow_ if supported only via workflow-derived mappings. The KE gene set score was then defined as the average per-gene confidence across all unique genes in the KE gene set:

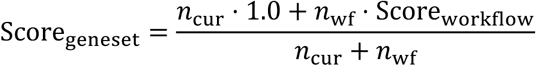

where n_cur_ describes genes supported by curated single-gene evidence and nwf genes supported only via workflow-derived mappings. Gene set scores ranged from 0.1 to 1 and reflected the relative contribution of curated versus workflow-derived evidence.

### 2.8 Gene/protein identification from KE titles

To capture explicit gene/protein/family mentions embedded in KE titles that may not be represented by pathway/ontology mappings, KE titles were additionally processed using Gilda for biomedical named entity recognition and grounding (Gyori et al., 2022). Dictionary-based string matching using HGNC symbols and synonyms was evaluated but found unreliable due to ambiguity and synonym overlap; therefore, Gilda was used to identify candidate text spans and link them to standardized entities. Input consisted of only KE titles. The entity types for genes (HGNC), proteins (UniProt), and protein families/complexes (FamPlex) were selected for gene retrieval. Candidates with grounding score ≥0.5 were retained and subsequently manually reviewed to remove spurious matches and confirm correct grounding. Curated grounded entities were used to augment KE gene sets, particularly for KEs referring to specific genes or proteins, where pathway- or ontology-based annotation was often not feasible or insufficiently specific. Genes added through this procedure were explicitly labeled as derived from FamPlex, HCGNC or UniProt in the final annotation.

### 2.9 AOP-level hazard class and regulatory endpoint categorization

To support a more regulatory-relevant interpretation of the AOP framework, AO/AOPs were mapped to regulatory hazard endpoint classes using a predefined endpoint vocabulary spanning Classification, Labeling, and Packaging (CLP) health hazards, CLP environmental hazards, the new CLP endocrine disruption hazard classes (ED HH/ED ENV; 2023/707), OECD emerging health hazards, and OECD/REACH terrestrial ecotoxicology (non-CLP) endpoints (plant, soil invertebrate, pollinator, bird, soil microorganism/STP inhibition). Neurotoxicity, immunotoxicity and developmental neurotoxicity were retained as separate OECD emerging health hazard annotation categories to preserve mechanistic and biological resolution, rather than being collapsed into STOT or reproductive toxicity. Endpoints were assigned according to the regulatory domain and biological interpretation of the AO: human health AOs were mapped to CLP health hazards (e.g., carcinogenicity, germ cell mutagenicity, STOT RE/SE) and/or OECD emerging health hazards when the AO reflects functional impairment in those domains, consistent with CLP’s health-hazard framing of target-organ toxicity as adverse functional effects not covered by other health hazard classes. Environmental AOs were mapped to CLP Hazardous to the aquatic environment when effects pertain to aquatic organisms (including algae/aquatic plants as part of aquatic hazard assessment) and to ED ENV where an endocrine mode of action plausibly drives adverse effects in environmental organisms, reflecting CLP’s separation of endocrine disruption hazards for humans versus the environment. Terrestrial-specific effects were mapped to OECD/REACH terrestrial ecotoxicology endpoint classes (plants, soil invertebrates, pollinators, birds, soil microorganisms/STP) when the AOP evidence or applicability indicated these organism groups. Taxonomic applicability information from AOP-Wiki was used, where available, as supporting evidence for assigning the regulatory domain of each AOP/AO. However, because taxonomic applicability was sometimes missing, incomplete, overly broad, or not fully aligned with the regulatory interpretation of the AO, endpoint assignments were not based on taxonomy alone. In such cases, biological plausibility, the identity of the AOP KEs, organism-specific biology, and the intended regulatory meaning of the endpoint class were considered. Thus, AOPs were mapped to human-health or environmental endpoints based on the combined interpretation of taxonomic applicability, mechanistic plausibility, and adverse outcome context. AOPs with evidence restricted to rats or mice were not automatically considered human-relevant. For these AOPs, human relevance was evaluated based on the biological plausibility of cross-species transferability of the underlying mode of action, distinguishing rodent-specific tumour mechanisms or apical readouts from pathways in which upstream molecular, cellular, or endocrine perturbations are conserved across mammals. Where human relevance was uncertain or limited, AOPs were annotated with “not assigned - taxonomic AD outside of mapping scope”. No endpoint was assigned when an AO represented a non-adverse pharmacological effect (e.g., analgesia) or when evidence did not support any class. Multi-label assignments were allowed.

### 2.10 AO-to-MeSH mapping for disease and literature context

To support disease-oriented interpretation and literature retrieval, AOs were mapped to MeSH descriptors using a semantic similarity-based approach. AO titles and MeSH term names were embedded in a common vector space with the all-mpnet-base-v2 sentence-embedding model (Reimers & Gurevych, 2019; Song et al., 2020) and SapBert (Liu et al., 2020), and candidate MeSH terms were ranked for each AO using cosine similarity for both models. The top 5 scoring candidates of both models were then manually curated to define the final MeSH annotations, allowing retention of multiple complementary MeSH terms where appropriate. Manual expert evaluations provided additional MeSH annotations in some cases.

### 2.11 CTD-derived AOP fingerprint

To demonstrate practical use of the KE-gene set annotation in a chemical prioritization setting, we reproduced the reference chemical groups introduced by Saarimaki et al. (2023a) by taking the chemical lists directly from their Supplementary File 1 (hepatotoxicants, thyroid-active chemicals, sex hormone receptor agonists (SHR), and carcinogens). For each chemical, we then constructed a chemical-associated gene set using CTD curated chemical-gene/protein interactions retrieved from the CTD bulk file CTD_chem_gene_ixns (downloaded March 31, 2026, 13:30:48 EDT), which provides curated interactions together with organism labels, NCBI GeneIDs, and CAS identifiers where available. AOP fingerprints were computed following the same general strategy as Saarimaki et al. (2023a). Briefly, for each chemical-associated gene set, we performed enrichment against AOP-level gene sets (defined as the union of genes across KEs within an AOP) and KE-level gene sets. Enrichment was performed using a one-sided Fisher’s exact test with Benjamini-Hochberg correction. Significant enrichments were summarized as an AOP “fingerprint” by ranking enriched AOPs/KEs by adjusted p-value and reporting top N results (as in Saarimaki et al. (2023a)).

## 3. Results and discussion

### 3.1. Semantic retrieval and LLM-assisted refinement improve KE mapping performance

We developed an AI-assisted, multi-step annotation workflow to generate an updated KE-to-gene mapping across all KEs represented in AOP-Wiki at the time of annotation, improving both coverage and precision compared with previous efforts (Saarimaki et al., 2023a). In contrast to the single-step NLP-based annotation used by Saarimaki et al. (2023a), our approach is multi-step and leverages (i) Sentence-BERT’s (SBERT) ability to retrieve semantically related ontology/pathway terms at high recall and (ii) an LLM’s ability to perform meaning-oriented filtering of those candidates (Figure 1).

**Figure 1:**
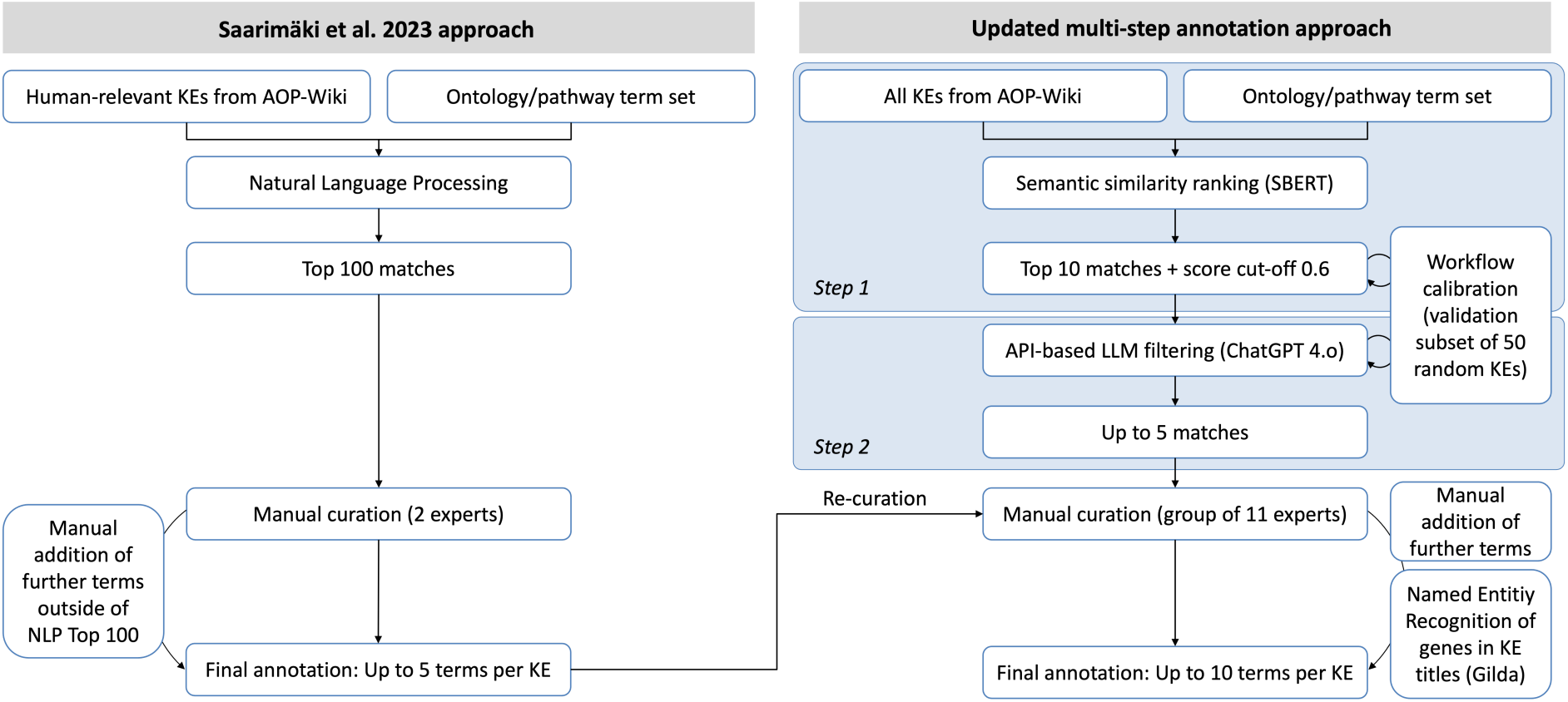
Comparison of annotation approaches between Saarimäki et al. 2023 and the updated annotation.

In the first stage, KE titles are matched to pathway and ontology term names using SBERT, retrieving the top candidates per KE (Fig. 1). SBERT generates semantically meaningful vector representations of short texts, known as embeddings, allowing KE titles and ontology/pathway terms to be compared based on meaning rather than exact word overlap, for example using cosine similarity (Reimers & Gurevych, 2019). Because AOP-Wiki KE titles are typically short but semantically dense, we hypothesized that embedding-based retrieval would capture paraphrase-level similarity more robustly than earlier NLP-based lexical matching approaches.

To quantify and optimize the first step, we calibrated the SBERT configuration (including the candidate subset forwarded downstream) using a manually curated validation subset of 50 randomly chosen KEs, labeled as true/false KE-term matches. On this validation set, SBERT candidate generation achieves precision = 0.26 and recall = 0.75, consistent with high coverage but low exactness (Tab. 1). This suggests that SBERT retrieval effectively captures semantically related candidates but requires further refinement to remove mechanistically inaccurate candidates.

**Table 1:** Retrieval performance of ontology-term matching methods for 2023 NLP: Saarimäki et al. 2023 NLP ranked predictions (Top100), 2026 Step 1 - SBERT: Novel annotation approach’s SBERT candidate retrieval (Top10), and 2026 Step 2 - LLM: Novel annotation approach’s LLM-filtered selections from SBERT candidates. Metrics were computed per KE and averaged across evaluable KEs. k: cutoff rank; metrics computed using only the top k predicted terms per KE. Precision@k = (# correct among top k) / (# returned among top k), for the LLM (C), which outputs variable-length lists up to 5, the denominator is the number of returned predictions up to k. Recall@k = (# correct among top k) / (# curated correct terms for that KE). Hit@k: fraction of KEs with ≥1 correct term in the top k. Blank entries for A, B at k 100 reflect that for SBERT/LLM only outputs up to a given rank were used, whereas the NLP pipeline from Saarimäki et al. 2023 used Top 100.

| k | Precision |  |  | Recall |  |  | Hit |  |  |
| --- | --- | --- | --- | --- | --- | --- | --- | --- | --- |
|  | 2023 NLP | 2026 Step 1: SBERT | 2026 Step 2: SBERT + LLM | 2023 NLP | 2026 Step 1: SBERT | 2026 Step 2: SBERT + LLM | 2023 NLP | 2026 Step 1: SBERT | 2026 Step 2: SBERT + LLM |
| 1 | 0.429 | 0.568 | 0.724 | 0.152 | 0.322 | 0.414 | 0.429 | 0.568 | 0.724 |
| 10 | 0.153 | 0.294 | 0.558 | 0.474 | 0.838 | 0.817 | 0.627 | 0.889 | 0.879 |
| 100 | 0.017 |  |  | 0.544 |  |  | 0.688 |  |  |

In the second step, the top scoring SBERT candidates are submitted to the API-accessed LLM ChatGPT 4.o to determine whether each candidate term truly matches the KE title in meaning and mechanistic intent, and to retain only the most appropriate mappings (up to five, Fig. 1). This LLM filtering step yields strong performance on the same curated validation subset, achieving hit@5 = 0.97, indicating that the correct term was almost always retained within the final set of up to five candidates. Importantly, the 50 KE validation subset was used not only to assess technical performance, but also to refine the LLM prompt and generation settings. This process is critical because LLM outputs are well documented to be sensitive to prompt formulation, including meaning-preserving variations in formatting and instruction phrasing that can lead to substantial differences in measured performance (Chen et al., 2025; He et al., 2024). Dedicated frameworks for analyzing prompt sensitivity demonstrate that LLM robustness varies across tasks and datasets, and that instance-level prompt variation can materially affect both objective performance and subjective evaluations, underscoring the need for systematic prompt assessment (Zhuo et al., 2024). Notably, the 50 KE subset used during workflow refinement did not fully reflect the final annotation standard, because final expert curation applied a more refined curation and consolidation logic than the initial development stage. To quantify the performance under the final annotation standard, we therefore evaluated retrieval performance against the final expert-curated reference annotations and compared these results to the Saarimaki et al. (2023a) annotations (Tab. 1). To disentangle the contributions of the individual components, we specifically compare (i) the SBERT retrieval step alone, (ii) the full SBERT + LLM pipeline, and (iii) the earlier NLP-based approach of Saarimaki et al. (2023a). This enables us to separately quantify improvements arising from semantic retrieval, and from the combined multi-step workflow based on SBERT and subsequent LLM filtering. Across all metrics, performance improves markedly from the SBERT candidate retrieval step to the LLM filtering step, with the most pronounced improvements observed in precision. This pattern indicates that the LLM step is effectively acting as a semantic verification layer that removes incorrect or only loosely related candidates. Related work in information retrieval and biomedical normalization similarly emphasizes that downstream reranking or refinement can substantially improve the quality of candidate sets produced by an initial high-recall retrieval step (Borchert et al., 2024). We also observe a small decrease in recall and Hit@k after LLM filtering, suggesting that in rare cases the LLM rejects candidates that human experts ultimately considered acceptable. This trade-off is expected when moving from a retrieval-oriented stage (designed for coverage) to a stricter semantic filtering stage (designed for precision).

Both Step 1 SBERT retrieval and Step 2 SBERT+LLM filtering achieve consistently higher performance across all reported metrics than the earlier Saarimaki et al. (2023a) NLP-based approach (Tab. 1). Importantly, this advantage was observed even though the 2023 approach used a much larger candidate list (Top 100), whereas our workflow used only the Top 10 SBERT-ranked candidates for downstream refinement. This indicates that SBERT retrieval already provides stronger semantic matching than lexical/NLP-style ranking alone, and that subsequent LLM filtering further improves precision. This is consistent with findings from other biomedical mapping tasks showing that embedding-based semantic approaches substantially outperform string-based approaches for ontology/term mapping, particularly when terminology varies and concepts are expressed heterogeneously (Liu et al., 2023).

Although direct comparison across studies is limited by differences in task definition and evaluation setup, the Step 2 LLM-refined outputs are broadly consistent with reported performance patterns in related ontology/entity mapping tasks, where lexical baselines tend to perform substantially worse than semantic retrieval methods, and an additional reranking or refinement step can further improve final selection (Borchert et al., 2024; Liu et al., 2023; Son Do et al., 2024).

Because the final annotation was determined through expert group curation, differences in the number of annotated terms per KE reflect curator decisions about which candidate terms were sufficiently mechanistically appropriate to retain. Under this final curation, the number of annotated terms per KE is lower in the updated annotation with roughly one-third fewer terms per KE compared to Saarimaki et al. (2023a) (Fig. 2A), indicating a more selective final mapping. This reduction is not driven by the automatic retrieval step alone, but by expert decisions to exclude or assign lower confidence to terms judged less appropriate representations of the KE. Notably, this reduction in terms per KE does not translate into increased gene-level specificity as measured by the number of KEs per gene (Supplementary file 1, Fig. S1). This suggests that annotation selectivity at the term level does not automatically yield a more discriminative gene-to-KE annotation, likely reflecting that many genes are multifunctional and pleiotropic, contributing to multiple processes, tissues, and phenotypes, so their recurrence across KEs is expected rather than necessarily indicative of nonspecific annotation (Barbitoff et al., 2025; Pritykin et al., 2015). Importantly, such gene recurrence does not diminish their relevance for any given KE, but instead reflects the shared molecular architecture underlying many biological and toxicological processes (Espinosa-Cantu et al., 2020).

**Figure 2:**
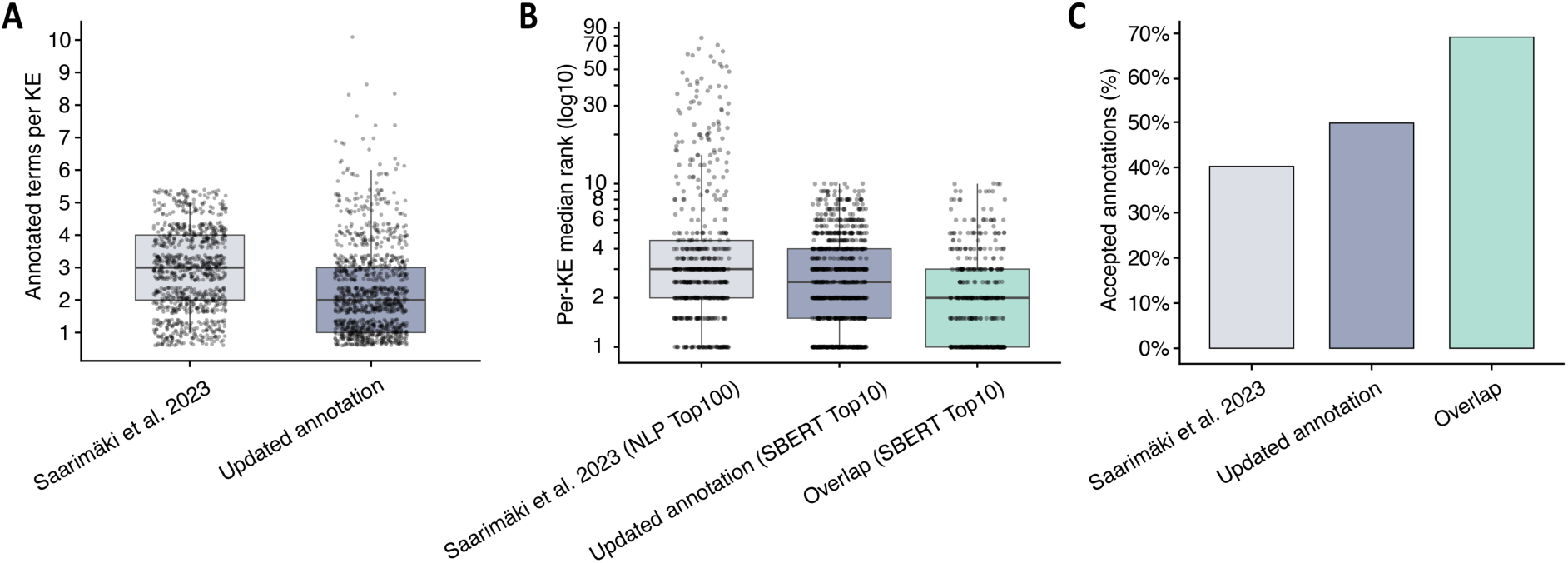
Annotation density, retrieval rank, and curation acceptance across annotation versions. (A) Distribution of the number of annotated terms per KE for Saarimäki et al. (2023) and the novel annotation. Boxplots summarize the median and interquartile range; points represent individual KEs. (B) Distribution of per KE median retrieval ranks based on score (log10 scale; lower is better) for three settings: Saarimäki et al. (2023) NLP Top 100, novel annotation approach SBERT Top 10, and overlapping KEs/terms evaluated with SBERT Top 10. In each workflow, candidate terms were ranked by their respective retrieval score (NLP-based matching score in Saarimäki et al. (2023); cosine similarity in the SBERT workflow), and each point represents the median rank of the accepted terms for one KE. (C) Curation acceptance rate (percentage of candidate annotations accepted) for Saarimäki et al. (2023), the novel annotation, and the overlap subset. Overlap refers to KEs/terms present in both annotation sets (shared subset), and ranks are computed with respect to each method’s corresponding candidate list (Top 100 for NLP, Top 10 for SBERT).

Because the final expert curation re-curated and integrated mappings from Saarimaki et al. (2023a), we next compare how candidate annotations from each source were treated during expert group curation. Overall, candidates proposed by the novel multi-step workflow are accepted more frequently (50%) than those originating from the Saarimäki et al. set (40%), while candidates assigned by both approaches show the highest acceptance (70%, Fig. 2C). Both the Saarimaki et al. (2023a) NLP-based matching and the Step 1 SBERT-based matching generate ranked candidate KE-term lists, in which higher-ranked (closer to rank 1) candidates received stronger matching scores from the respective retrieval method. Interestingly, overlapping candidates also tend to be ranked more favorably, i.e., they have closer to 1 (better) mean ranks than candidates unique to either approach (Fig. 2B). This pattern is informative in two aspects. First, it suggests that the novel multi-step pipeline produces candidates that align more consistently with curator expectations for mechanistic plausibility. This is particularly notable given that these Saarimäki et al. annotations already contained extensive manual curation and augmentation (50% of terms annotated manually), whereas the updated annotation candidate pool was not manually curated or augmented at the candidate-generation stage and required very few curator-added terms during final curation (≈2% of the final annotation). This suggests that the updated annotation workflow produces candidate annotations that are already more often judged biologically meaningful by expert curators, thereby reducing the need for manual augmentation during curation substantially. Second, the elevated acceptance of overlapping terms (Fig. 2C) suggests that agreement between independent retrieval strategies provides a practical robustness signal, consistent with information retrieval data-fusion findings that consensus across systems correlates with relevance (“chorus effect”) (Lillis, 2020).

### 3.2. Expanded KE coverage and biologically coherent gene sets in the updated annotation

To assess the practical utility of the updated annotation, we first quantify coverage across KEs and AOPs. The AI-assisted, multi-step annotation approach produced gene set representations for 1,254 KEs, which together span 523 AOPs, leaving only 7 AOPs without KE-level gene set coverage (Fig. 3). Compared with the previous annotation by Saarimaki et al. (2023a), we annotated 385 additional KEs, while 52 previously proposed KE annotations were rejected during re-curation (Supplementary file 1, Tab. S1) and 49 KEs from the prior set were no longer present in the current AOP-Wiki snapshot (Supplementary file 1, Tab. S1). To extend coverage beyond pathway- and ontology-based mappings, we added a complementary annotation stream that captures explicit gene/protein/family mentions embedded directly in KE titles. Such KEs often refer to a specific molecular target (e.g., receptor, transcription factor, enzyme, complex), where pathway-level gene sets can be too diffused or where no suitable ontology term exists that is both specific and mechanistically aligned. This procedure produced 463 candidate grounded entities after score filtering, of which 398 were confirmed by manual review, resulting in curated single-gene (or gene-family/protein) identifications across 335 KEs. Together with retained title-derived gene annotations from Saarimaki et al. (2023a), a total of 340 KEs were ultimately supported by title-derived gene evidence. Of these, 112 KEs were supported exclusively by title-derived gene evidence, as no pathway/ontology term annotation was ultimately accepted for those KEs (Fig. 3). Importantly, we track gene provenance (HGNC, UniProt, FamPlex) explicitly, allowing users to distinguish title-derived identifications from pathway/ontology-derived gene sets and to interpret or filter these evidence types separately when appropriate.

**Figure 3:**
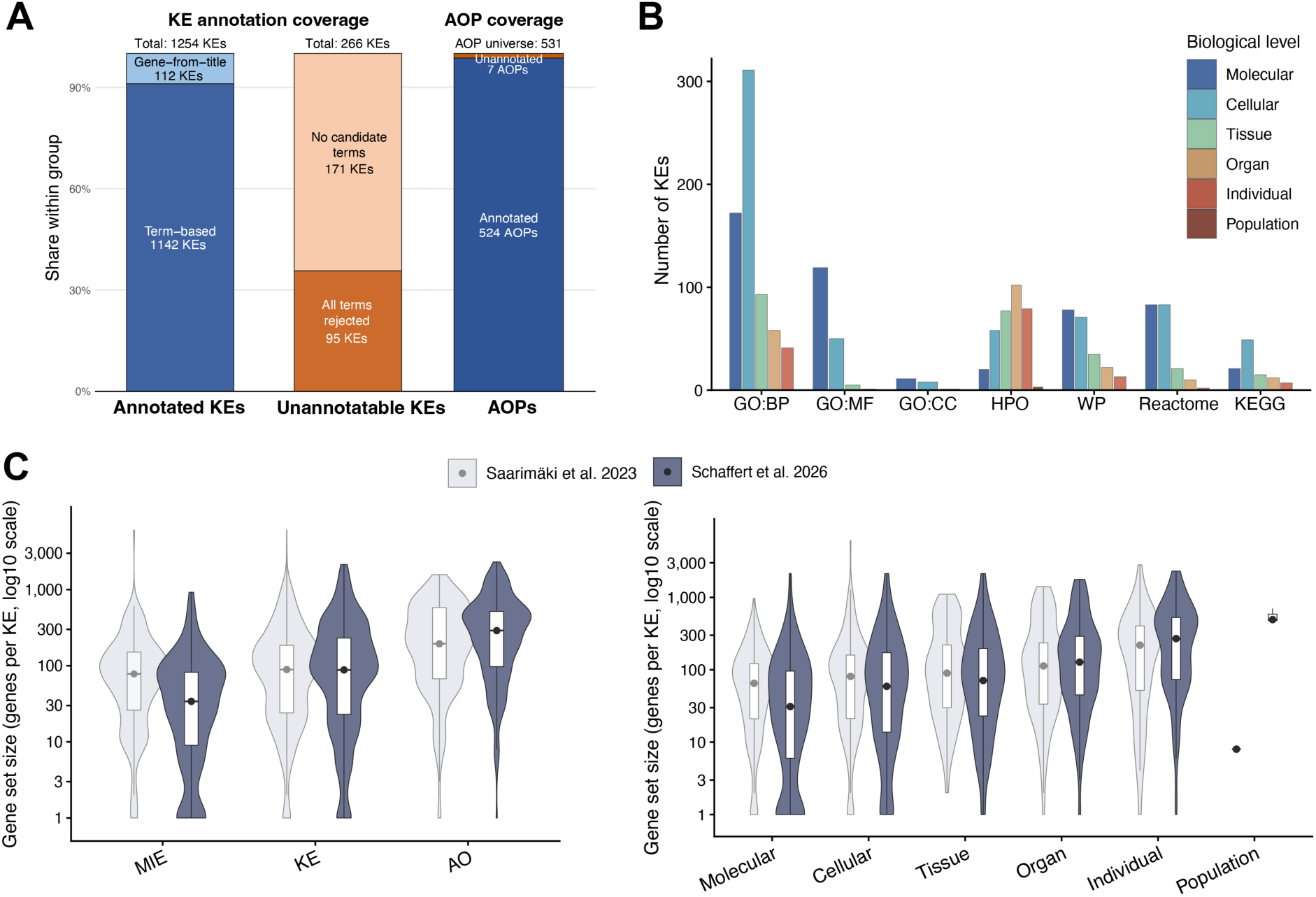
Coverage and characteristics of KE gene set annotations. (A) Number of AOPs covered by at least one annotated KE versus AOPs without coverage and counts of KEs, showing annotated KEs by Saarimäki et al. 2023, annotated KEs by the present updated annotation, and non-annotatable KE. (B) Number of KEs annotated per knowledgebase source (GO, HPO, WikiPathways, Reactome, KEGG) stratified by biological level. (C) Distribution of gene set sizes (genes per KE; log10 scale) across event categories (MIE/KE/AO) and KE biological levels.

Despite this broad coverage, 266 KEs remained unannotated (Fig. 3). Of these, 171 received no candidate pathway/ontology terms or title-derived single-gene annotations from either the novel annotation workflow or the re-curated Saarimaki et al. (2023a) annotation. The remaining 95 KEs had such candidate term annotations, but all were rejected during expert curation. In most cases, these represent events that are currently difficult to translate into gene sets in a biologically meaningful way, for example because the KE description is (i) too unspecific (e.g., *Event:1239 Altered, Gene Expression, Event:2007 Non-coding RNA expression profile alteration, Event:2302 Increase, Malformations*) (ii) framed at a level not captured by the used pathway/ontology terms (e.g., *Event:1036 Increased, liposarcoma, Event:2233 Decreased, ERαβ heterodimers, Event:245 Activation, PXR/SXR*), or (iii) not sufficiently mechanistically resolved to support a gene-based representation (e.g., *Event:1550 Deposition of Ionizing Energy; Event:961 Increased, Clearance of thyroxine from serum; Event:947 Increase, Early Life Stage Mortality, Event:829 Damage, Lipid bilayer*). In other words, these KEs highlight the present boundary of what can be “annotated” into gene sets using existing knowledgebases and controlled vocabularies.

Across biological levels reported in the AOP-Wiki, KEs differ systematically in the types of resources that support their gene annotations. GO terms contribute the largest share to molecular and cellular KEs, with declining contributions toward tissue/organ/individual/population levels and a similar gradient is observed for curated pathway resources (Reactome, WikiPathways and KEGG) (Fig. 3B). This pattern is biologically coherent because lower-level KEs often describe discrete molecular activities or proximal cellular processes that align naturally with mechanistic ontologies and pathway models. In line with GO’s organization, GO Molecular Function annotations are concentrated at molecular/cellular levels, consistent with the prevalence of MIE-like KEs describing receptor, enzyme or transcription factor binding and activity changes, whereas GO Cellular Component becomes particularly appropriate when KEs are fundamentally about localization or translocation events. When KEs describe broader processes or states, such as activation of a pathway, coordinated stress responses or changes in cell state, GO Biological Process and curated pathway databases provide a more appropriate representation than molecular-function terms, because they encode multi-step or multi-component biology more directly. In contrast, HPO contributes proportionally more to higher-level KEs, which matches its design as an ontology of phenotypic abnormalities and therefore its closer alignment with integrated organ- and organism-level events than with narrowly mechanistic molecular descriptions. This shift from GO/pathway annotations at lower levels to HPO at higher levels is consistent with ontology structure and biological organization, supporting appropriate KE granularity and mapping plausibility. Although this alignment was not explicitly imposed in the automated workflow, it emerged naturally from the meaning-aware annotation and subsequent curation. Based on this pattern, future KE-to-gene annotation efforts, particularly when extending the workflow to additional KEs or AOPs, may benefit from aligning the biological level and mechanistic scope of the KE with the resource type best suited to represent it: GO Molecular Function for direct molecular activities, GO Cellular Component for localization/translocation, GO Biological Process and pathway knowledgebases for process-level effects, and phenotype ontologies like HPO for higher-level outcomes. This alignment helps preserve mechanistic intent, reduces category mismatches (e.g., forcing higher-level outcomes into overly granular mechanistic terms), and provides an additional plausibility check for mapping quality when the dominant resource type matches the KE’s biological scale.

Across the annotated KEs, gene set size shows a clear upward shift with increasing biological level (molecular, cellular, tissue, organ, individual, population) (Fig. 3C). A similar pattern is observed for MIE, KE, AO distributions (Fig. 3C), which often corresponds to these biological levels of organization. This trend is biologically plausible: higher-level phenotypes and outcomes integrate perturbations across multiple pathways, cell types, and regulatory circuits, whereas lower-level molecular events are typically mediated by more constrained molecular events, aligning with a systems and network biology perspective in which higher-order phenotypes emerge from interactions across multiple biological scales. Consequently, higher-level KEs may be better represented by broader, integrative gene sets reflecting coordinated activity across pathways, cell types, and cellular systems, whereas lower-level molecular KEs are more often captured by smaller and more mechanistically constrained gene sets (Fischer et al., 2025; Parikshak et al., 2015). Importantly, this biological hierarchy is more pronounced in the present annotation than in the previous approach, driven by smaller gene set sizes at lower biological levels, especially molecular and cellular KEs (Fig. 3C). This pattern is consistent with improved annotation robustness at the lower biological levels, where narrowly defined events are less likely to be inflated by overly broad matches.

A further improvement is the disappearance of previously extreme outlier gene sets, e.g., 1259 *Event: Narcosis* and *Event:169 Disruption of membrane integrity*, previously mapped to very large gene sets, including ∼4,000 genes for membrane disruption, which are not observed at comparable magnitudes in the current annotation (Fig. 3C, Supplement 1 Fig. S2). This likely reflects reduced susceptibility to generic term annotation under surface-level matching. In the updated workflow, clearly inconsistent terms for narcosis (GOCC plasma membrane, GOBP regulation of plasma membrane organization) were discarded by the curators, and ambiguous membrane-related terms (e.g., GOCC plasma membrane, GOBP membrane disruption in other organism) were replaced by more mechanistically appropriate process terms identified by the LM-approach (GOBP regulation of plasma membrane repair, GOBP regulation of membrane permeability). Large gene sets do remain, over 30 KEs exceed 1,000 genes, but these are predominantly higher-level events (organ/individual level), where broadness is expected because such KEs summarize integrated physiology rather than discrete molecular mechanisms. One prominent exception occurs for broadly defined cellular inflammation KEs (e.g., Increase/Increased, inflammation), which exceed ∼1,800 genes. In these cases, multiple ontology/pathway terms from GO, WikiPathways, and HPO match well by name and definition, but because “inflammation” is itself a highly general biological event, the union of these matched terms spans a very broad gene space. This breadth is consistent with the biology of inflammation, which is a core response that can be triggered by diverse insults including toxic compounds, and it encompasses multiple pathways, mediators, and cell types (e.g., cytokine signaling, inflammasome activation, immune recruitment) (Chen et al., 2018). However, the magnitude of this effect compared to other KEs highlights a practical gap in the current AOP-Wiki event space: inflammation-related KEs are often under-granular, collapsing distinct inflammatory processes into a single broad KE label. Given the centrality of inflammation in toxicant responses and downstream pathology, increased granularity (e.g., separating acute vs chronic inflammation, innate vs adaptive components, inflammasome-driven vs cytokine-dominant mechanisms, tissue-specific inflammatory contexts) would likely improve mechanistic resolution and reduce ambiguity in KE-level gene set interpretation (Bender et al., 2025; Chen et al., 2018).

Comparing KE-specific gene memberships between the previous and current releases reveals substantial divergence among KEs shared with Saarimaki et al. (2023a). Using the Jaccard index (JI) to quantify overlap, 58% of shared KEs showed low similarity at the gene set level (JI < 0.3) (Supplementary file 1; Fig. S3). In contrast, similarity was higher at the level of annotated term identifiers, where 40% of KEs had JI < 0.3. Together, these results indicate that gene set divergence is often driven by changes in gene membership that can occur even when term-level mappings are comparatively stable, but also that term-level changes account for a substantial fraction of the observed gene set differences.

To understand the drivers of these changes, we traced gene set differences back to the provenance of term assignments and term availability across knowledgebase releases. In most cases, changes in gene set composition co-occurred with changes in annotated term identifiers (i.e., terms added and/or removed) (Supplementary file 1, Tab. S2). We observed two prominent, conceptually distinct causes behind term-level changes. First, a considerable fraction of KEs was affected by terms that were introduced into the MSigDB-based term universe over time, meaning that such terms could not have been proposed or recovered under the older Saarimäki MSigDB term set and were therefore legitimately absent from the earlier annotation. Second, some previously used terms are now missing from the current MSigDB-derived term universe and therefore could not be matched by the current workflow (“missing in term KB”). Where appropriate, these terms were retained through re-curation of the earlier annotation.

Importantly, we also identified a small subset of KEs where gene sets changed despite unchanged term identifiers (five KEs; Supplementary file 1, Tab. S2). This pattern is consistent with updates in the underlying biological knowledgebases themselves: resources such as GO, Reactome, WikiPathways, and HPO are actively maintained, and the gene membership associated with a stable identifier may change across releases as curation evolves. In practical terms, this means that version-to-version differences in KE gene sets can reflect both (i) updated KE-term mapping decisions and (ii) changes in the gene membership of retained term identifiers, which can occur as the underlying resources are updated. The latter suggests a practical route for future maintenance, because gene set content could in principle be refreshed automatically for stable curated term mappings without repeating manual annotation. This underscores the importance of explicitly reporting knowledgebase provenance (release/version) and treating KE gene sets as time-stamped representations of curated biological knowledge.

Inspection of KEs with low Jaccard similarity between the updated annotations and Saarimaki et al. (2023a) annotations revealed a consistent pattern (Supplementary file 1, Tab. S3). Generic or weakly aligned terms from Saarimaki et al. (2023a) were often replaced by more mechanistically specific and context-appropriate terms, shifting from broad cellular responses or umbrella pathways toward disease-, process-, or receptor subtype-level concepts. For example, *Event:1291 Hepatotoxicity* was previously represented by the highly general GO biological process term cellular response to toxic substance, which describes a broad stress response rather than hepatotoxicity pathogenesis. In the updated annotation, this was replaced by more specific phenotype-level terms capturing clinically and mechanistically meaningful hepatic outcomes (e.g., hepatic failure and hepatic necrosis). Similarly, *Event:1138 Uncoupling of oxidative phosphorylation, reduced ability to generate ATP* moved from broad oxidative phosphorylation pathways toward terms explicitly capturing uncoupling as the core mechanistic concept (including mitochondrial uncoupling-specific entries). In *Event:1038 Activation, beta2 adrenergic receptor*, a more generic “adrenergic receptor binding” term was refined to a beta2-specific molecular function annotation, improving subtype specificity. Likewise, downstream reproductive and pathology-related KEs show a systematic shift toward phenotype-aligned specificity: *Event:1515 Spermatocyte depletion* moved from generic apoptosis-related processes to terms explicitly referencing spermatogenesis and spermatocyte maturation arrest; and *Event:1763 Urothelial tumor* shifted from broad cancer-related terms to cancer type-specific annotations (e.g., bladder/urinary tract neoplasm and bladder cancer pathways). In addition to increased specificity, several examples indicate removal of broad or ambiguously matched terms that likely inflated gene sets without adding mechanistic resolution. For *Event:1511 Lipid peroxidation*, the updated annotation explicitly captures the lipid oxidation component of the event, while dropping less targeted membrane-related terms. *Event:1547 Mitochondrial injury* also illustrates improved alignment: a loosely associated “mitochondrial uncoupling” mapping was replaced by annotations more consistent with mitochondrial dysfunction and apoptosis-related mitochondrial changes.

Several low-overlap examples reflect not only revised mapping decisions but also changes in the available term universe over time. For instance, *Event:1044 Promotion, mesovarian leiomyomas* was previously approximated using loosely related smooth muscle growth/proliferation terms, whereas the updated annotation assigns the direct phenotype term uterine leiomyoma, which was not available in the earlier input term catalog and therefore could not have been selected previously.

Moreover, 56 KEs previously annotated in Saarimaki et al. (2023a) were rejected during re-curation, providing informative examples of mismatches introduced by lexical matching (Supplementary file 1, Tab. S3). Several rejections reflect token-level confounding, for example, *Event:1686 Deposition of Energy* which refers to physical energy deposition by a stressor (e.g., radiation), was mapped to mechanistically unrelated energy metabolism/homeostasis terms. Likewise, *Event:1913 Endothelial cell dysfunction* was mapped to the GO term for endothelial cell activation, and *Event:1421 Activated, LXR* was linked to the PPAR signaling pathway, suggesting manual supplementation in the absence of suitable lexical candidates by the NLP-based approach of Saarimaki et al. (2023a). In these cases, the updated approach assigned low confidence and did not retain the mappings, consistent with improved semantic specificity.

Together, these changes suggest that low gene set overlap frequently reflects meaningful refinement, shifting from broad, loosely associated representations toward terms that more directly encode the KE’s mechanistic intent, and, in some cases, the removal of previously assigned mappings that expert re-curation judged to be mechanistically mismatched.

### 3.3. Expert-derived confidence scores expose annotation uncertainty

Rather than treating KE-gene set annotations as uniformly reliable, we encoded expert judgement as an explicit confidence layer attached to each KE mapping, implemented through double-independent expert review and rule-based consolidation. Curators provided qualitative assessments (fit, partial, not_fit), enabling derivation of a quantitative quality score for each KE annotation. This score provides an explicit confidence layer that reflects curator agreement and annotation specificity and can be propagated to the gene set level. In practice, this enables confidence-aware downstream analyses, such as AOP fingerprinting, where users can identify, weight, or even filter KE gene sets based on annotation quality, improving transparency and interpretability compared to unscored gene set resources.

Across all scored KEs, 57% of KEs achieved high scores (≥ 0.7), and an additional 31% fell in the intermediate range (0.3-0.7), indicating that for the majority of KEs, the final retained mappings were judged by curators to be biologically relevant (Fig. 4A). At the same time, a smaller subset showed substantial uncertainty, including 12% with low scores (< 0.3). These low-score cases are informative because they identify KEs for which available ontology/pathway terms do not provide a well-aligned representation, or for which the KE itself is intrinsically difficult to capture as a gene set.

**Figure 4:**
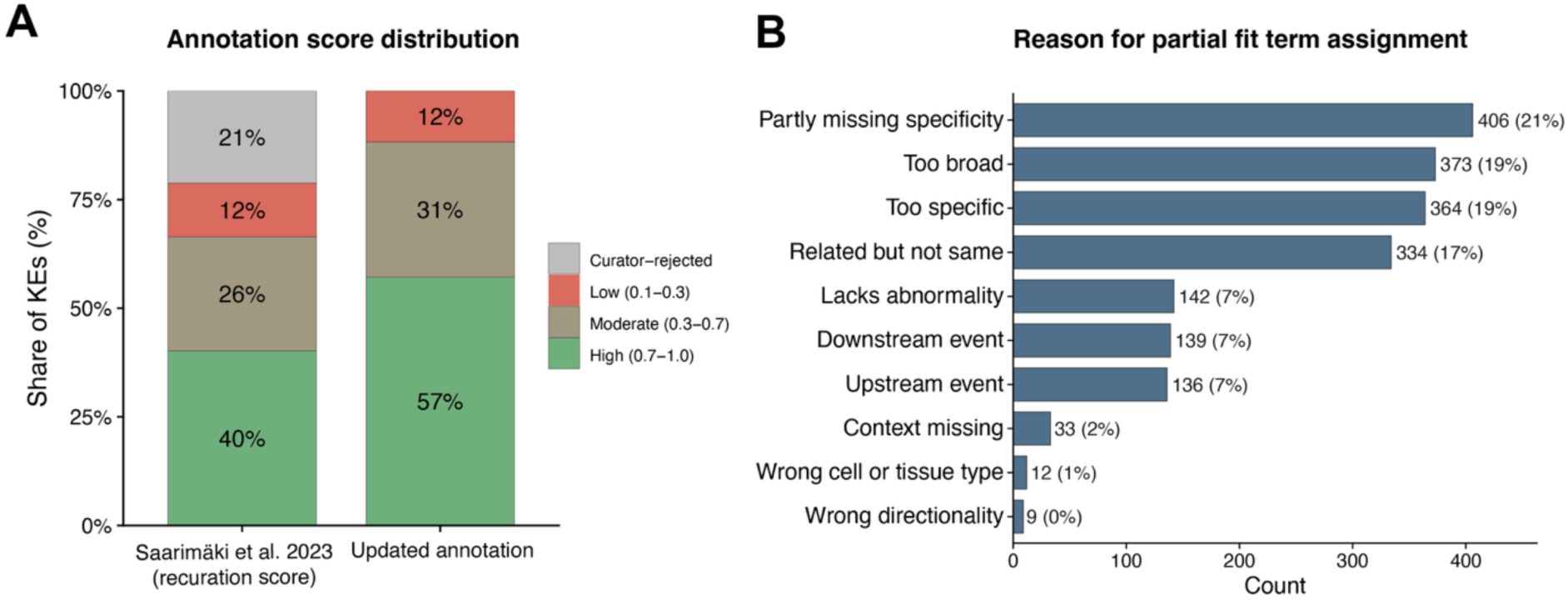
Curation-derived annotation confidence scores and reasons for uncertainty. (A) Distribution of KE annotation scores (range 0.1–1.0) for Saarimäki et al. 2023 annotations (re-curated) and full updated annotation set, binned into three intervals with the corresponding KE counts shown above each bar. (B) Most frequent curator reason codes contributing to partial KE-term inclusions, highlighting common sources of uncertainty.

As the previously published Saarimaki et al. (2023a) annotations were re-curated within the same expert-review framework used for the present study (Fig. 1), this enables a direct comparison of KE-term curator assessment. Under this common scoring scheme, the earlier annotation was rated systematically lower than the final updated annotation: only 40% of KEs achieved a high score when based on Saarimäki-derived mappings, and 21% of KEs had no retained mappings after re-curation (i.e., all previously assigned mappings for that KE were rejected). Together, these results indicate that the updated workflow yields final KE mappings that are more frequently endorsed by expert review.

To understand sources of uncertainty among retained mappings, we analyzed curator reason codes assigned to partial-fit inclusions (Supplementary file 1, Tab. 1; Fig. 4B). These codes captured why a candidate term is biologically related but still insufficient to represent the KE as written. The most frequent reasons are partly_missing_specificity (21%), too_broad (19%), too_specific (19%), and related_but_not_same (17%), followed by lacks_abnormality (7%) and causal-placement mismatches (downstream_event 7%, upstream_event 7%). Collectively, these patterns show that low scores are most often driven by granularity mismatch (terms broader or narrower than the KE) and by conceptual proximity without full equivalence (related but not the same).

Across reasons for exclusions (rejected candidate terms, summarized separately in Supplementary file 1, Fig. S4), too_broad and related_but_not_same are again prominent, indicating that many candidates were biologically adjacent but still not acceptable as KE representations at the level required for gene set modeling.

The confidence score differentiates KEs that are straightforward to represent from KEs that remain difficult to annotate (Tab. 2). High-scoring KEs (Tab. 2; score = 1.0) typically correspond to events for which well-aligned terms exist in the knowledgebases used in this study, and curators therefore converge on clear mappings (e.g., estrogen receptor activity, PPARα activation, adipogenesis, programmed cell death). In contrast, low-scoring KEs (Tab. 2; score = 0.1) tend to fall into recurring categories: (i) very specific or context-dependent events (e.g., life-stage or disease-context specificity such as infant leukemia, pancreatic acinar cell tumors, granulosa cell proliferation of gonadotropin-independent follicles), (ii) highly granular molecular statements for which no close ontology/pathway term was available in our sources (e.g., AHR/ARNT dimerization), (iii) events that are hard to represent as gene sets overall (e.g., physiologic readouts such as decreased mucosal blood flow or membrane integrity disruption), and (iv) composite KE phrasing that combines multiple concepts into a single KE title (e.g., systemic inflammation leading to hepatic steatosis), which makes it difficult to find a single term that matches the whole statement.

**Table 2:** Examples for KEs with top (score = 1.0) and with the lowest annotation scores (score = 0.1).

| KE ID | KE name | Annotation score | Reason for decreased score, annotated term is: |
| --- | --- | --- | --- |
| Event:149 | Increase, Inflammation | 1 | - |
| Event:1060 | Alteration, lipid metabolism | 1 | - |
| Event:1864 | Increase, Programmed cell death | 1 | - |
| Event:1029 | Increased, adipogenesis | 1 | - |
| Event:1395 | Liver Cancer | 1 | - |
| Event:1276 | Lung fibrosis | 1 | - |
| Event:227 | Activation, PPARα | 1 | - |
| Event:1541 | Mitochondrial Complex I inhibition | 1 | - |
| Event:948 | reduced production, VEGF | 1 | - |
| Event:1606 | Activation of hepatic stellate cells | 1 | - |
| Event:748 | Increased, Estrogen receptor (ER) activity | 1 | - |
| Event:193 | Decreased, Nitric Oxide | 0.1 | downstream_event |
| Event:1254 | Infant leukaemia | 0.1 | partly_missing_specificity, related_but_not_same |
| Event:177 | Mitochondrial dysfunction | 0.1 | too_specific, related_but_not_same |
| Event:944 | dimerization, AHR/ARNT | 0.1 | partly_missing_specificity |
| Event:1270 | Inactivation of PPARγ | 0.11 | too_broad, partly_missing_specificity |
| Event:1800 | Granulosa cell proliferation of gonadotropin-independent follicles, Reduced | 0.1 | too_broad, partly_missing_specificity |
| Event:1404 | Decreased, mucosal blood flow | 0.1 | too_broad, partly_missing_specificity |
| Event:486 | systemic inflammation leading to hepatic steatosis | 0.1 | partly_missing_specificity, related_but_not_same |
| Event:2005 | Altered neurotransmission in development | 0.1 | partly_missing_specificity, related_but_not_same, too_broad |
| Event:169 | Disruption, Membrane integrity | 0.1 | lacks_abnormality, related_but_not_same, downstream_event |
| Event:1725 | Pancreatic acinar cell tumors | 0.1 | too_broad, partly_missing_specificity, context_missing |

Two examples highlight different “resolution limits” of ontology/pathway-based representation. First, mitochondrial dysfunction is a central toxicology concept but is often used as an umbrella label; within the pathway/ontology sources used here, no single term captured the concept at the resolution implied by the KE, and candidate matches were either too specific or only related but not equivalent. Second, the contrast between PPARα and PPARγ shows the opposite situation: the KE is mechanistically precise, but the knowledgebases available for mapping in this study do not provide terms at the required resolution. Activation, PPARα received a perfect score because PPARα-specific pathway/ontology terms were available and could be matched directly. In contrast, PPARγ-related KEs scored low because our knowledgebases did not provide sufficiently PPARγ-specific pathway/ontology terms; the closest candidates were broader PPAR signaling terms that cover multiple isoforms. This matters biologically because different PPARs regulate overlapping but distinct gene processes and have distinct functions, so a broad PPAR pathway mapping for a PPARγ-specific KE should be interpreted cautiously (Wagner & Wagner, 2020).

The distribution of partial-fit and rejection reasons points to improvement opportunities across three layers. 1) AOP-Wiki/KE development: frequent “too broad”, “partly missing specificity”, and “context missing” cases suggest that some KEs are under-specified or combine multiple concepts. Clearer separation of causal steps, explicit context qualifiers, and more consistent KE phrasing would improve interoperability. The OECD AOP Developers’ Handbook already emphasizes structured KE description and transparent evidence assembly (OECD, 2018), stronger recommendations for ontology-aware KE phrasing could further improve machine-actionability and interoperability (Mortensen et al., 2025).

2) Ontologies/pathways: cases such as missing isoform-level receptor biology indicate that controlled vocabularies can lack resolution for some toxicology-relevant mechanisms, motivating targeted pathway curation and expansion, including community-driven resources such as WikiPathways that support ongoing content growth and community contribution (Agrawal et al., 2024). 3) AI-assisted workflow: the partial-fit and rejection patterns show that automated matching alone is not sufficient at AOP-Wiki scale. KE titles often encode mechanistic and other contextual constraints that are not reliably captured from term names alone. Resolving these cases requires toxicology and AOP-specific understanding, so expert review remains necessary. Thus, we conclude that large-scale KE gene set validity cannot be assessed objectively without human judgment.

These results also highlight the practical value of the AI-assisted, multi-step annotation design. Automation is effective for generating a manageable candidate selection at scale, while structured expert review provides the final mechanistic decision and the accompanying confidence score layer. The annotation score can support weighted and transparent evidence summaries, which is often what is needed in regulatory-relevant contexts (OECD, 2021). Specifically, the score can be used to distinguish between high-certainty KE mappings that can be emphasized as robust mechanistic evidence and lower-certainty mappings that can still be reported but clearly flagged as more ambiguous or context-dependent. This makes the AOP fingerprint easier to communicate as a structured summary of mechanistic evidence, while remaining explicit about uncertainty and provenance. Such confidence-aware reporting aligns with OECD guidance that emphasizes structured and transparent evidence assembly in AOP development and evaluation (OECD, 2018, 2021), and it supports broader community efforts toward machine-actionable and FAIR-oriented AOP resources where quality signals and provenance metadata facilitate responsible reuse (Mortensen et al., 2025). In the same spirit, it complements ongoing OECD efforts to improve the transparency and reproducibility of omics evidence used in regulatory settings, where reporting frameworks aim to standardize how omics analyses are documented and interpreted (OECD, 2023b).

Overall, the score is not only a confidence indicator for AOP/KE enrichment analyses, but also a diagnostic signal that identifies KEs whose current wording is difficult to translate into gene sets without added context or decomposition and areas where pathway/ontology resources may need more targeted terms to represent toxicology-relevant mechanisms.

### 3.4. Regulatory endpoint and MeSH mapping place annotated AOPs in a broad toxicological and disease context

In addition to the KE-level gene set annotation, we assigned AOP-level contextual descriptors to support interpretation of downstream analyses. Specifically, we mapped AOP adverse outcomes to regulatory hazard/endpoint classes and assigned MeSH descriptors for disease-oriented and literature-oriented contextualization. These mappings were generated at the AOP/AO level and are therefore conceptually distinct from the KE-level gene set annotations described above. The regulatory endpoint mapping provides a structured summary of the annotated AOP landscape in terms of toxicological and regulatory relevance. Viewed across the full annotated AOP space, the regulatory endpoint mapping indicates that the current AOP landscape spans a broad set of toxicologically relevant domains. Among human-relevant categories, the largest numbers of AOPs map to STOT SE/RE, followed by endocrine disruption – human health, toxicity for reproduction, and carcinogenicity, while substantial numbers also cover emerging or increasingly prioritized regulatory domains such as adult neurotoxicity, developmental neurotoxicity (DNT), and immunotoxicity (Fig. 5). This breadth is consistent with the growing use of AOPs in regulatory toxicology and new approach methodologies across multiple endpoint domains, including endocrine disruption, neurotoxicity, developmental neurotoxicity, and immunotoxicity (European Food Safety et al., 2024; OECD, 2023a).

**Figure 5:**
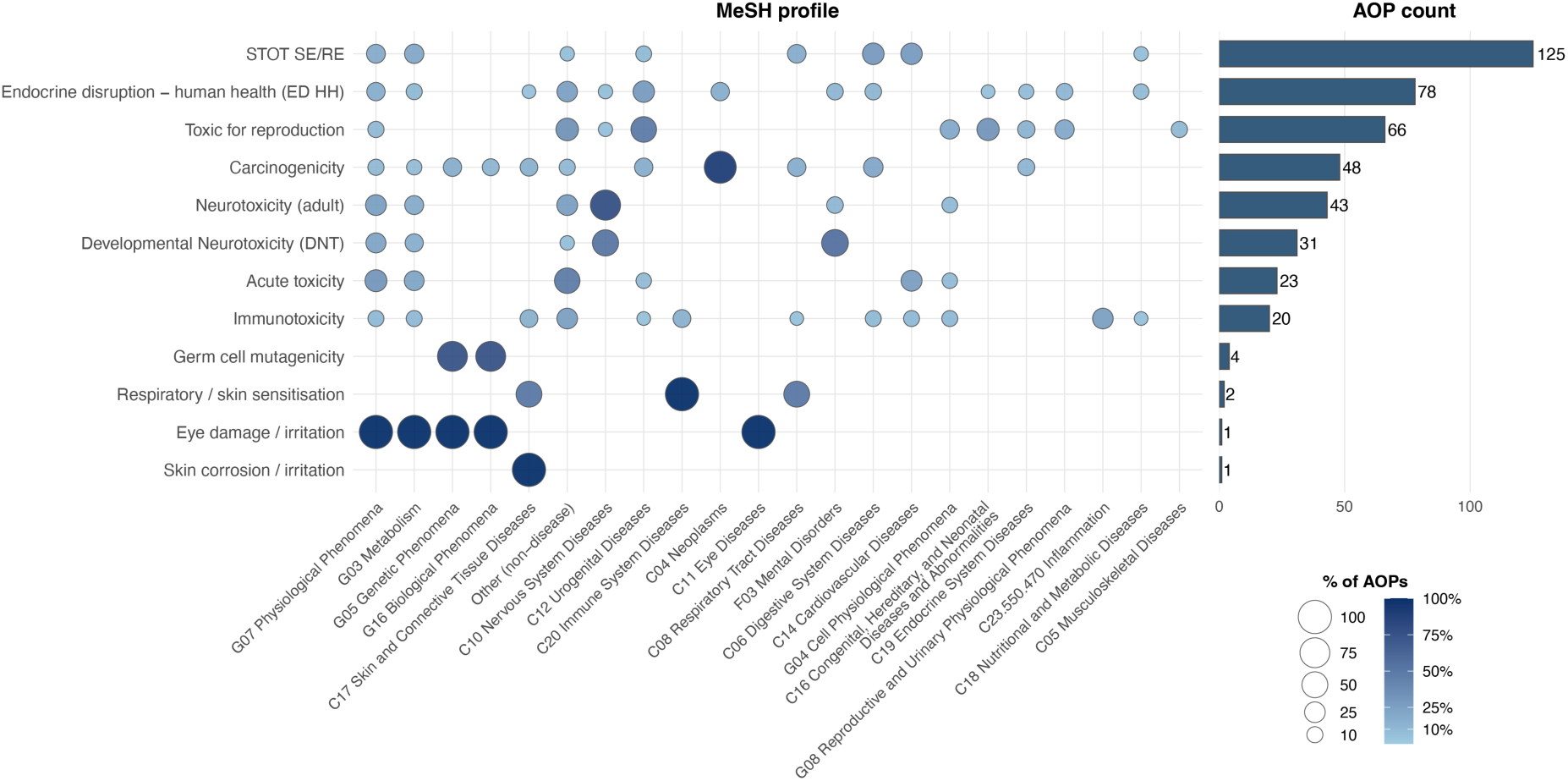
MeSH patterns and AOP counts across human-relevant hazard/regulatory endpoint classes. Dots show the percentage of AOPs in each hazard/regulatory endpoint that contains an AO mapped to the MeSH category. Bars show the count of AOPs per hazard/regulatory endpoint. Multiple assignments per AOP are possible. STOT SE/RE = Specific target organ toxicity single exposure/repeated exposure.

The MeSH assignments provide an additional disease- and phenotype-oriented view of these AOPs. Some endpoint classes show highly intuitive profiles: carcinogenicity is dominated by Neoplasms, toxic for reproduction is overrepresented for urogenital and congenital/hereditary disease categories, and adult neurotoxicity and DNT show strong representation of Nervous System Diseases, with DNT also showing a substantial contribution from Mental Disorders, consistent with the long-recognized developmental and functional consequences of perturbing neurodevelopmental processes (Pitzer et al., 2023). By contrast, STOT SE/RE and endocrine disruption – human health are associated with a more diverse MeSH space, which is biologically plausible because these hazard classes cut across multiple organ systems and disease areas rather than corresponding to a single disease family. The endocrine-disruption pattern is also compatible with the broad systemic roles of endocrine signaling across metabolism, development, reproduction, and tissue homeostasis. A notable pattern is that acute toxicity is enriched for non-disease or broader physiological MeSH categories. This is unsurprising, because adverse outcomes in AOPs are not restricted to formal diseases and often include functional or apical biological effects relevant to risk assessment. MeSH itself also contains hierarchical branches for physiological phenomena and other non-disease concepts, so mapping to such categories can be valid. However, the recurrence of “Other (non-disease)” categories across several endpoint classes also reflects that some AOP adverse outcomes are currently phrased in ways that are biologically meaningful but not harmonized with disease-oriented controlled vocabularies. This points to a broader opportunity for future AOP development: more systematic AO naming and explicit disease-oriented annotation could improve interoperability with external resources such as Comparative Toxicogenomics Database (CTD), which relies on controlled disease vocabularies integrating MeSH, and could strengthen literature linkage and regulatory interpretability.

Overall, the combined endpoint and MeSH analysis suggest that the current AOP landscape comprises a diverse and increasingly regulatory-relevant set of biological outcome domains. The KE-level gene set annotation developed here therefore supports not only mechanistic enrichment, but also interpretation across a broad spectrum of human-health and environmental endpoint contexts.

### 3.5. Expanded annotation supports confidence-aware and endpoint-informed interpretation of omics data in an AOP context

To demonstrate downstream utility of the newly developed annotations, we investigated the AOP fingerprint using the reference chemical groups defined previously (Saarimaki et al., 2023a): carcinogens, hepatotoxicants, thyroid-active chemicals, and sex hormone receptor agonists (SHR). Using curated CTD chemical-gene interactions as input, we performed enrichment analysis against gene sets defined at both the AOP and KE levels. Compared to the Saarimaki et al. (2023a) annotation, the present annotation yields a higher number of significantly enriched terms per chemical across all tested categories, with the increase being more pronounced at the AOP level than at the KE level (Fig. 6A). This is consistent with the increased KE coverage achieved by the novel approach, which expands the number of KE gene sets available to contribute evidence to AOP-level enrichment.

**Figure 6:**
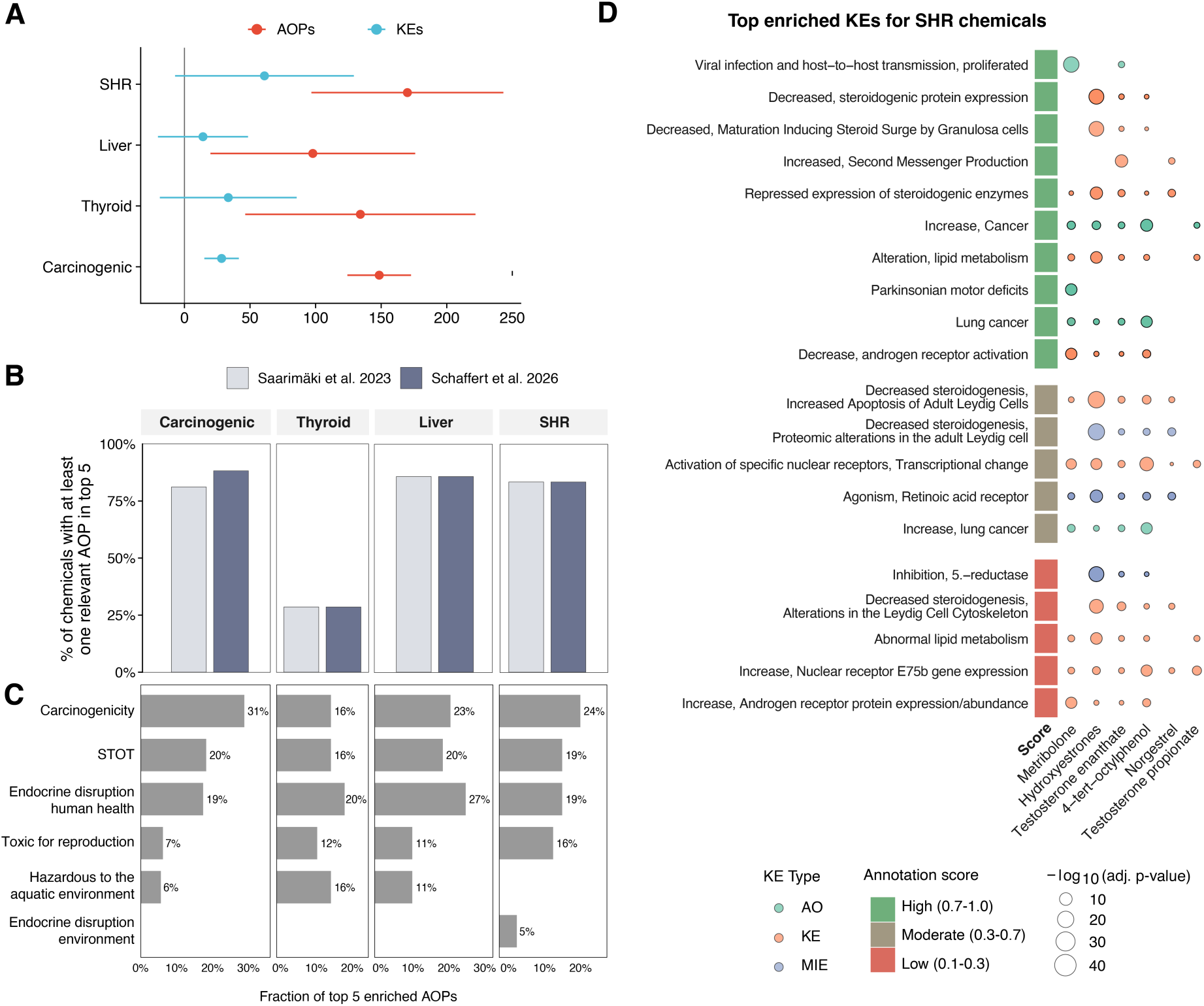
CTD-based AOP fingerprinting benchmark and confidence-aware interpretation. (A) Mean difference in the number of significantly enriched AOPs and KEs per chemical for each chemical group (Carcinogenic, Thyroid, Liver, SHR), computed as novel annotation − Saarimäki et al. 2023 (padj < 0.05). Points indicate the mean difference across chemicals, horizontal bars show 95% confidence intervals, left vertical line marks no change. (B) Fraction of CTD chemicals whose top 5 enriched results include at least one expected endpoint-relevant AOP, comparing the novel annotation to the Saarimäki 2023 annotation. (C) Regulatory endpoints and their distribution across the top 5 enriched AOPs for each chemical group using the new annotation. (D) Example KE-level fingerprint of the top 20 enriched KEs for the sex hormone receptor (SHR) group using the new annotation.

Despite this overall increase in enriched terms, the endpoint-level interpretability remains comparable or improves depending on the chemical class. When investigating if the top 5 contain at least one expected AOP, the updated annotation shows an improved fraction of correct hits for carcinogenic chemicals, while liver and SHR categories remain high and similar between annotations; thyroid chemicals remain challenging in both annotations (Fig. 6). The persistent difficulty for thyroid chemicals suggests that limited performance is not solely an annotation coverage issue, but also reflects limits in specificity of CTD-derived chemical-gene signatures for thyroid-active compounds and/or remaining gaps in mechanistic AOP/KE representation for thyroid-relevant biology, despite ongoing progress in developing and assembling thyroid AOP networks in recent years (Noyes et al., 2019; Wiklund et al., 2023).

To assess regulatory relevance beyond hit counts, we examined the hazard classes and regulatory endpoints associated with adverse outcomes of the top-ranked AOPs (Fig. 6C). For carcinogenic chemicals, the largest share of top 5-ranked AOPs is expectedly annotated with the hazard category of Carcinogenicity. Notably, STOT SE/RE and Endocrine disruption - human health (ED HH) also appear prominently among the top categories. Chemical carcinogenic outcomes frequently arise through cytotoxicity-induced regenerative proliferation and endocrine modulation, which promote sustained proliferative signaling and tumorigenic processes across multiple tissues (Cohen, 2024; Del Pup et al., 2016; Strupp et al., 2023). For the thyroid-active group, ED HH is among the dominant categories, which is coherent with the endocrine nature of thyroid hormone system disruption. This disruption can have an impact on multiple different organ systems (Noyes et al., 2019; Wiklund et al., 2023), explaining the prominent enrichment of STOT-related AOPs. The additional presence of aquatic/environmental hazard categories in this group is also mechanistically plausible because thyroid hormone signaling is highly conserved across vertebrates and thyroid-related AOP networks explicitly integrate endpoints spanning aquatic taxa (e.g., amphibian and fish developmental outcomes) (Haigis et al., 2023; Zwahlen et al., 2024). For liver-active chemicals, the enrichment profile is dominated by ED HH together with substantial STOT representation. This combination is biologically coherent because liver toxicity frequently involves endocrine-regulated metabolic processes and nuclear receptor modulation, including PPAR signaling, and many liver-relevant AOPs connect metabolic dysregulation, inflammation, and tissue remodeling to adverse outcomes such as steatosis, fibrosis, and tumor development (Monroy-Ramirez et al., 2021). Mechanistically, PPARs are central transcriptional regulators of hepatic lipid metabolism and inflammatory responses, and dysregulation of these processes is tightly coupled to liver disease that is frequently represented in AOP landscape (Monroy-Ramirez et al., 2021; Pawlak et al., 2015). From a hazard-class perspective, the appearance of STOT among top categories is consistent with the CLP systemic toxicity context that includes organ toxicity classifications. The SHR (sex hormone receptor agonist) chemical group shows a major share of top-ranked AOPs falling under Carcinogenicity and additional contributions from STOT, ED HH, and Toxic for reproduction. This pattern is mechanistically plausible because estrogen and androgen receptor signaling modulates transcriptional processes that are directly implicated in hormone-dependent cancers (most prominently breast and prostate cancer), and sex hormone signaling also underlies reproductive development and fertility endpoints (Kim et al., 2025). Taken together, top-ranked AOP fingerprints for each chemical group are connected to endpoint domains that are mechanistically aligned with the intended groups, while also showing plausible cross-domain connections (e.g., endocrine-metabolic-tumor interaction for liver and SHR).

Finally, we illustrate confidence-aware interpretation at the KE level using the SHR group (Fig. 6D). The top results include endocrine-relevant mechanisms, such as modulation of steroidogenesis and steroidogenic protein expression, 5α-reductase, and Leydig-cell steroidogenesis related events, which aligns well with the expected mechanisms of sex-hormone receptor agonists. The SHR fingerprint also contains a small set of low-confidence KEs (score ≤ 0.3), including *Inhibition 5α-reductase*, *Abnormal lipid metabolism*, and several KEs with narrow or ambiguous phrasing such as *Increase, Nuclear receptor E75b gene expression*, *Increase, Androgen receptor protein expression/abundance*, and *Decreased steroidogenesis*, *Alterations in the Leydig Cell Cytoskeleton*. These cases illustrate three recurring limitations already visible in the curation reason overview (Fig. 4B): Granularity mismatch (candidate terms are broader or narrower than the KE), related-but-not-equivalent matches, and context or phrasing constraints that are difficult to represent with the knowledgebases used for mapping in this study. Moderate-confidence KEs (0.3-0.7) include mechanistically plausible endocrine and transcriptional readouts such as *Activation of specific nuclear receptors*, *Transcriptional change*, *Agonism, Retinoic acid receptor*, and general tissue remodeling processes. These typically reflect situations where multiple pathway/ontology candidates are plausible but differ in scope or context, producing less curator consensus. In contrast, many endocrine-relevant KEs central to SHR mechanisms (e.g., androgen receptor activation/agonism and steroidogenesis-related events) are enriched with high confidence, indicating that for core mechanisms the mapping is both biologically aligned and curator-consistent.

### 3.6. Recommendations for use, interpretation and future extensions

The aim of this annotation is to provide, for each KE, a gene set representation that captures the molecular pathways and biological programs conceptually underlying the KE. Importantly, these gene sets are designed to be broadly applicable across experimental contexts, reflecting mechanistic content that may generalize across systems rather than being tailored to a single study design, tissue, or exposure scenario. As such, the resource is best viewed as a foundational mapping that enables AOP-informed analysis of transcriptomics (and other omics) data, supporting tasks such as mechanistic interpretation, KE-anchored enrichment analyses, and cross-system comparisons. Because KEs are used in diverse biological contexts, users may wish to refine mappings for specific applications. For example, in a targeted case study, one might restrict a KE gene set to genes expressed in a relevant cell type, modulate memberships based on tissue specificity, integrate exposure- or time-resolved expression patterns, or prioritize pathway branches that are most consistent with the AOP context. In other words, this annotation is intended to provide a mechanistically grounded starting point.

Critically, this resource is not intended to directly define biomarkers for testing or regulatory decision-making on its own. In particular, biomarkers are typically quantitative measures, whereas the present AOP-based annotation provides qualitative mechanistic associations and does not, at its current stage, specify the magnitude, direction, or threshold of change required for biomarker definition (FDA-NIH, 2016). Biomarker development therefore requires additional downstream steps, such as model-based feature selection, evaluation of predictive performance, and experimental confirmation, along with explicit consideration of confounders, assay constraints, and reproducibility (NRC, 2007). The value of the present annotation in that workflow is that it provides a basis for mechanistic hypothesis generation that can then be refined for biomarker selection using empirical data.

Several limitations should be considered when interpreting enrichment or AOP/KE fingerprints. The current implementation is restricted to a human gene space, even though annotated KEs participate in a broad range of AOPs. This means that KEs outside a strictly human toxicology context are currently represented through human-centered molecular annotations, which is useful for many conserved processes but should be interpreted accordingly. Pathway- and ontology-derived gene sets may also include genes associated with the broader process surrounding a KE, rather than only direct drivers of the event itself. Conversely, for KEs explicitly referring to individual genes or proteins, the incorporation of title-derived molecular entities improves specificity where pathway- or ontology-based matching alone would be insufficient.

Moreover, in most cases, the gene set representation does not encode the direction of a KE (increase/decrease; activation/inhibition). This reflects a general limitation of pathway/ontology gene sets, which typically define membership of genes associated with a process rather than the sign of regulation under a specific perturbation. Consequently, enrichment of a KE gene set should be interpreted as evidence that the molecular response involves genes linked to the KE’s biology, but not as direct evidence that the KE is increased vs decreased. Directionality, when needed, must be inferred from the omics data itself (e.g., sign of differential expression, signed enrichment statistics, or gene-level patterns). This remains non-trivial, because activation of a biological event may involve both up-and down-regulation across different components (e.g., inhibitors vs activators), similar to the interpretation challenges encountered in standard pathway enrichment workflows (Wang et al., 2024; Zhao & Rhee, 2023). This makes the annotation especially useful for comparative and integrative analyses, where the goal is to detect whether molecular profiles are consistent with the biology represented by that KE.

Even under the meaning-aware workflow presented here, some KEs remain challenging to translate into gene sets. This limitation reflects the boundary conditions of gene set representability given current controlled vocabularies and knowledgebases. In practice, KEs can be difficult to represent when they are (i) underspecified or highly context-dependent (e.g., missing relevant tissue/cell type or directionality), (ii) framed at a biological level not captured well by pathway/ontology terms, or (iii) not sufficiently mechanistically resolved to support a gene-based abstraction. Making these difficult-to-map KEs visible through our scored annotation is valuable, because it indicates where additional mechanistic clarification of the KE and improved ontology coverage may be required.

A promising direction for improving scalability and precision in the future may be to incorporate definitions (not only titles) into the annotation process. Sentence-level models can benefit from richer context, and pathway/ontology resources often provide concise definitions that describe the scope of the term. Integrating these definitions alongside term names could improve semantic alignment and reduce ambiguity during candidate retrieval and filtering. However, a key limitation on the KE side is that AOP-Wiki typically provides a KE description, but this text rarely provides a concise definition. These descriptions often include background information, related biology, examples, or narrative context, which dilutes the information needed for machine-actionable semantic matching. A simple but potentially high-impact improvement would be the addition of a concise “definition” field for KEs (and ideally AOPs) that captures the mechanistic essence in one or two sentences. Such structured definitions would not only support more robust language-model operability, but also improve interpretability, consistency, and FAIR reuse (Vogt et al., 2025). The observation that some KEs cannot be captured cleanly by existing ontologies suggests that such free text may be an essential complement to controlled vocabularies for representing complex biological events. Rather than treating free text as an obstacle, future AOP-Wiki evolution could embrace it as a structured resource: definitions and constrained textual fields can be operationalized via language models to improve annotation quality while acknowledging that not all mechanistic concepts are currently covered by formal ontologies (Lucy Lu Wang et al., 2018; Peng et al., 2023).

Overall, the annotation is best used as a confidence-aware framework for mechanistic interpretation. Its strongest value lies in enabling structured, transparent, and mechanistically broad interpretation of omics data across many KEs and AOPs at once, while preserving provenance and confidence information to guide context-appropriate use.

## Conclusion

In this work, we introduce a scalable AI-assisted, multi-step workflow for annotating AOP-Wiki KEs with mechanistically meaningful gene set representations, enabling AOP knowledge to be used directly in toxicogenomics. By combining semantic retrieval, LLM-assisted refinement, and structured expert group curation, the workflow translates heterogeneous KE phrasing into analysis-ready gene sets while attaching an explicit confidence layer grounded in expert judgment. This directly addresses a central obstacle for AOP-guided omics at scale: achieving broad coverage across an expanding KE landscape while making uncertainty transparent, particularly when KEs differ in granularity, context dependence, and feasibility of gene set representation.

A key contribution of this study is the demonstration that meaning-aware AI components outperform traditional lexical/NLP-based matching for KE-to-term mapping by producing candidates that are more consistently mechanistically plausible and more frequently accepted by experts. At the same time, the results reaffirm an essential point: AOP annotation fundamentally requires toxicological expert assessment to adjudicate mechanistic appropriateness, resolve ambiguity, and ensure that mappings remain biologically defensible. Importantly, our workflow substantially reduces manual curation effort by narrowing the candidate space and increasing candidate quality, thereby shifting expert time from exhaustive search to high-value review and consolidation while improving the overall annotation quality.

To maximize reuse and enable continuous improvement, we release the resulting annotation resource and associated metadata (including confidence scores, curator reason codes, and provenance). The workflow is explicitly designed to support routine updates, allowing the resource to evolve with AOP-Wiki growth and with changing pathway/ontology knowledgebases. Beyond immediate utility for KE-and AOP-level enrichment and fingerprinting, the approach provides a general, reusable blueprint for future AOP annotation efforts. In particular, it establishes a practical foundation for the OECD Omics2AOP (workshop in proof-stage at Archives of Toxicology), where the same multi-step approach and scoring concepts can be extended beyond transcriptomics to additional molecular layers (e.g., proteins and metabolites) and translated into community tools that support transparent, confidence-aware AOP-guided interpretation. Together, these contributions strengthen the connection between mechanistic AOP knowledge, multi-omics data integration, and regulatory-relevant evidence generation.

## Competing Interests

The authors declare no competing interests.

## Acknowledgements

This study was funded by the Horizon Europe Framework research and innovation programme (HORIZON), INSIGHT (grant number 101137742), and the European Research Council (ERC) programme, Consolidator project ARCHIMEDES (grant number 101043848). A. Schaffert, L. Möbus, and G. del Giudice were funded by the Tampere Institute of Advanced Study (IAS). The work of M. Paparella at the Medical University of Innsbruck is funded by the Austrian Federal Ministry for Climate Action, Environment, Energy, Mobility, Innovation and Technology, Department V/5 – Chemicals Policy and Biocides.

## Notes

### Competing Interest Statement

The authors have declared no competing interest.

