## Supplementary file 1 for "Integrating Semantic Retrieval, LLM-based Refinement, and Structured Expert Curation for Scalable AOP Gene Mapping"

### Supplement 1

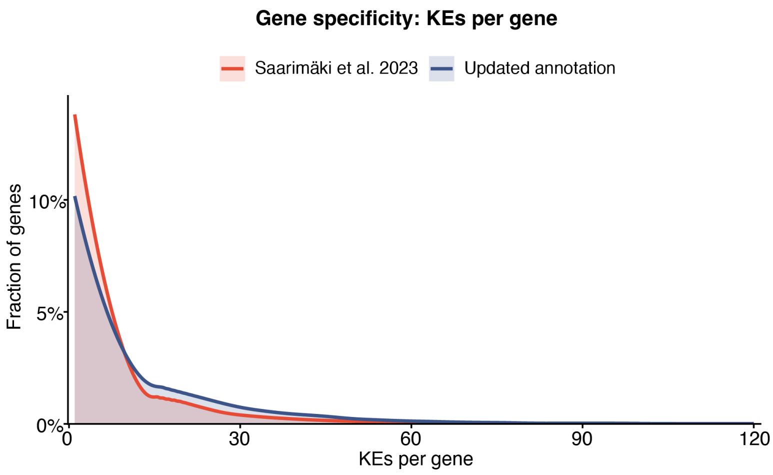


*Figure S1: Gene specificity represented as KEs per gene.*

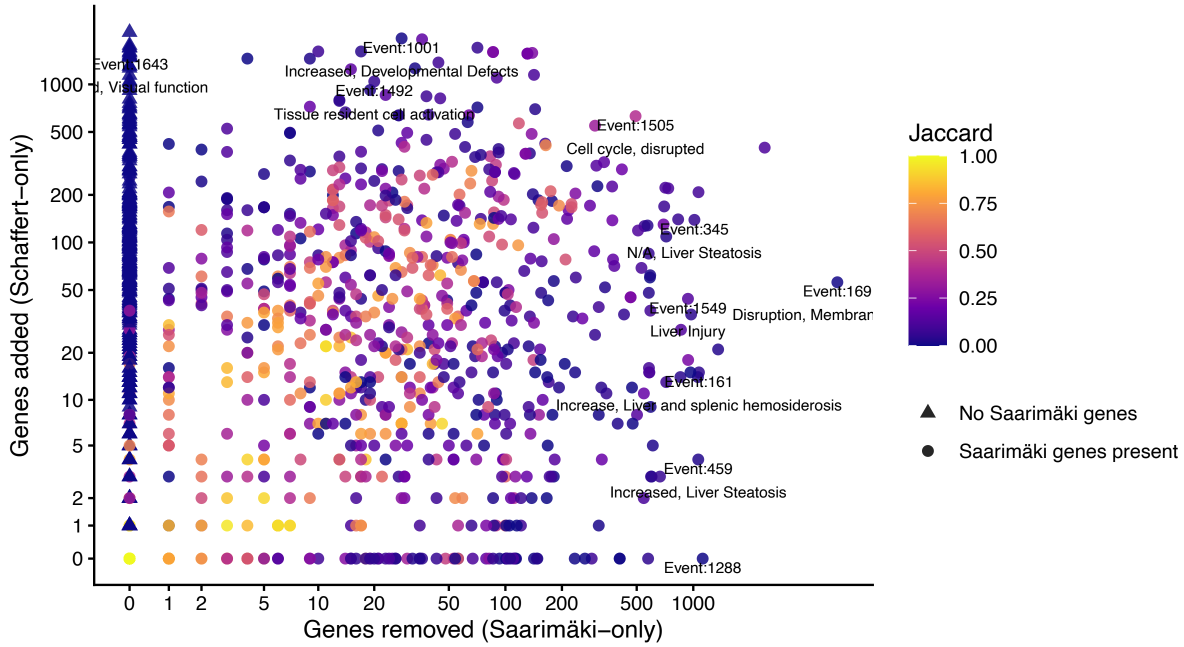


*Figure S2: Gene-set changes per KE.*

*Table S1: Overview of annotation coverage in numbers.*

| | Number |
|---|---|
| Input KE universe | 1503 |
| KEs with candidate annotation after Step 1 | 1236 |
| KEs with candidate annotation after Step 2 | 1099 |
| KEs receiving curated single-gene identifications | 335 |
| Additional KEs receiving curated single-gene identifications from Saarimäki 2023 annotation | 5 |
| Newly annotated KEs vs Saarimäki 2023 | 385 |

| | |
|---|---|
| Annotated KEs removed vs Saarimäki 2023 | 52 |
| Additional KEs removed due to deletion in AOP-Wiki | 49 |

*Table S2: Predefined reason codes for partial and not_fit annotations of KE-term candidates.*

| Category | Meaning |
|---|---|
| too_broad | Term covers a broad umbrella concept; KE is narrower/more specific. |
| too_specific | Term describes a narrow sub-process/subtype that does not represent the KE as written. |
| wrong_biological_process | Represents a different biological process than the KE. |
| wrong_directionality | Direction implied by the KE is not correctly captured by the term (e.g., KE is “decreased signaling” but term is “positive regulation”). |
| partly_missing_specificity | Term is in the right general area but lacks a key required component implied by the KE (not just “broad”, but incomplete). |
| context_missing | Term lacks required context (cell type, tissue, temporal stage) explicitly part of the KE. |
| wrong_cell_or_tissue_type | KE specifies a cell type/tissue/organ context, but the term is not specific to that context (e.g., generic/neutral), so the KE cannot be represented with the required anatomical resolution. |
| not_related | No conceptual relationship; term is irrelevant. |
| related_but_not_same | Term is clearly related but represents a different concept than the KE. |
| downstream_event | Term describes a consequence of the KE (a later event in the causal chain), not the KE itself. |
| upstream_event | Term describes a cause/preceding event that can lead to the KE, not the KE itself. |
| lacks_abnormality | Term describes a normal/physiological process while the KE requires abnormality/perturbation. |

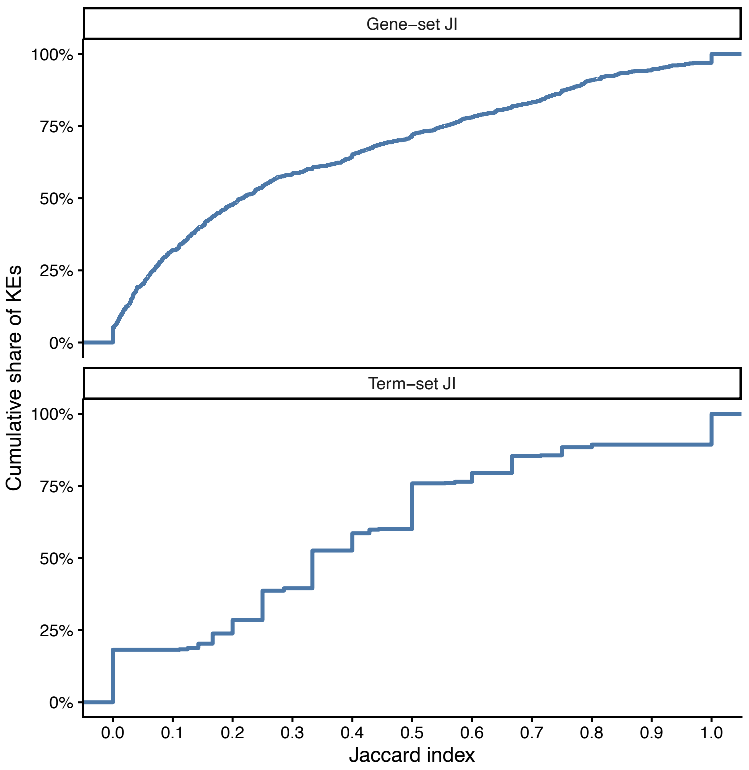


*Figure S3: Cumulative distributions of similarity between Saarimäki et al. (2023) and the updated annotation shared KEs. Empirical cumulative distribution functions (ECDFs) of Jaccard indices (JI) computed per key event (KE) for gene sets (top panel) and annotation term sets (bottom panel). For each KE, JI was calculated as the size of the intersection divided by the size of the union between the two approaches (range: 0–1), where 1 indicates identical sets and 0 indicates no overlap. The ECDF shows the cumulative fraction of KEs with JI ≤ x. KEs lacking a comparable set on either side (i.e., no genes or no terms available for one approach) were excluded from the corresponding panel. Higher ECDF values at low JI indicate a larger proportion of highly divergent KEs, whereas curves shifted toward higher JI indicate overall stronger agreement between Saarimäki and the updated annotation.*

*Table S2: Changes in KE gene sets in the updated annotation relative to Saarimäki et al. (2023), stratified by term-change categories and gene set agreement. Counts and percentages are reported relative to the total number of KEs in the updated annotation. Term-change categories describe whether term identifier sets were unchanged ("terms same") or differed due to KE-specific term additions/removals, terms unavailable in the current workflow candidate term universe ("missing in current term KB"), or terms newly available in the MSigDB-derived term universe in the current workflow ("MSigDB added over time"). Gene set agreement is assessed by set equality of Ensembl gene identifiers per KE. KE counts across term-change categories are not mutually exclusive; a KE may be counted in multiple categories if multiple term-change types occur within the same KE.*

| **KE gene set** | genes same | genes different | genes different | genes different | genes same | genes different | genes same | genes same | genes different |
|---|---|---|---|---|---|---|---|---|---|
| **term category** | terms same | | added to KE | removed from KE | missing in term KB | | removed from KE | added to term KB over time | |
| **number of KEs affected** | 5 | 88 | 635 | 552 | 1 | 251 | 2 | 2 | 290 |
| **% of total KEs** | 0,4 | 7,05 | 50,84 | 44,2 | 0,08 | 20,1 | 0,16 | 0,16 | 23,22 |

*Table S3: Selection of KEs with strongest difference in gene sets between the updated annotation and Saarimäki et al. (2023) annotations. KEs are displayed with the added, removed or still shared gene numbers, and the corresponding added terms or removed Saarimäki et al. terms.*

| **KE_ID KE_name** | **added** | **removed** | **shared** | **Updated annotation added terms** | **Saarimaki removed terms** |
|---|---|---|---|---|---|
| Event:1291 Hepatotoxicity | 199 | 12 | 0 | HP_ACUTE_HEPATIC_FAILURE [added_to_KE]<br>HP_CHRONIC_HEPATIC_FAILURE [added_to_KE]<br>HP_HEPATIC_FAILURE [added_to_KE]<br>HP_HEPATIC_NECROSIS [added_to_KE] | GOBP_CELLULAR_RESPONSE_TO_TOXIC_SUBSTANCE [removed_from_KE] |
| Event:1138 Uncoupling of oxidative phosphorylation, Reduced ability to generate ATP | 8 | 178 | 0 | GOMF_OXIDATIVE_PHOSPHORYLATION_UNCOUPLER_ACTIVITY [MSigDB_added_over_time]<br>REACTOME_MITOCHONDRIAL_UNCOUPLING [added_to_KE] | GOBP_NEGATIVE_REGULATION_OF_OXIDATIVE_PHOSPHORYLATION [missing_in_current_KB]<br>GOBP_OXIDATIVE_PHOSPHORYLATION [removed_from_KE]<br>GOBP_REGULATION_OF_OXIDATIVE_PHOSPHORYLATION [removed_from_KE]<br>KEGG_OXIDATIVE_PHOSPHORYLATION [removed_from_KE] |
| Event:1044 Promotion, mesovarian leiomyomas | 10 | 90 | 0 | HP_UTERINE_LEIOMYOMA [MSigDB_added_over_time] | GOBP_MUSCLE_HYPERTROPHY [removed_from_KE]<br>GOBP_POSITIVE_REGULATION_OF_SMOOTH_MUSCLE_CELL_PROLIFERATION [removed_from_KE]<br>GOBP_SMOOTH_MUSCLE_CELL_PROLIFERATION [removed_from_KE] |
| Event:1038 Activation, beta-2 adrenergic receptor | 16 | 1 | 0 | GOMF_BETA_2_ADRENERGIC_RECEPTOR_BINDING [added_to_KE]<br>2. BETA_2_ADRENERGIC_RECEPTOR [other_added] | GOMF_ADRENERGIC_RECEPTOR_BINDING [removed_from_KE] |
| Event:1511 Lipid Peroxidation | 84 | 70 | 0 | GOBP_LIPID_OXIDATION [added_to_KE] | WP_OXIDATIVE_DAMAGE [missing_in_current_KB]<br>GOBP_MEMBRANE_LIPID_CATABOLIC_PROCESS [removed_from_KE] |
| Event:1515 Spermatocyte depletion | 207 | 25 | 0 | HP_SPERMATOCYTE_MATURATION_ARREST [MSigDB_added_over_time]<br>HP_SPERMATOGENESIS_MATURATION_ARREST [MSigDB_added_over_time]<br>HP_ABNORMAL_SPERMATOGENESIS [added_to_KE] | GOBP_EXECUTION_PHASE_OF_APOPTOSIS [removed_from_KE] |
| Event:1547 Mitochondrial Injury | 59 | 6 | 0 | GOBP_APOPTOTIC_MITOCHONDRIAL_CHANGES [added_to_KE]<br>HP_MITOCHONDRIAL_RESPIRATORY_CHAIN_DEFECTS [added_to_KE] | REACTOME_MITOCHONDRIAL_UNCOUPLING [removed_from_KE] |

| | | | | | |
|---|---|---|---|---|---|
| Event:1763 Urothelial Tumor | 188 | 3 | 0 | HP_BLADDER_NEOPLASM [added_to_KE]<br>HP_URINARY_TRACT_NEOPLASM [added_to_KE]<br>KEGG_BLADDER_CANCER [added_to_KE]<br>WP_BLADDER_CANCER [added_to_KE] | GOBP_UROTHELIAL_CELL_PROLIFERATION [missing_in_current_KB]<br>GOBP_RESPONSE_TO_TUMOR_CELL [removed_from_KE] |
| Event:716 Increase, cell proliferation (hepatocytes) | 17 | 207 | 0 | GOBP_POSITIVE_REGULATION_OF_HEPATOCYTE_PROLIFERATION [added_to_KE]<br>GOBP_REGULATION_OF_HEPATOCYTE_PROLIFERATION [added_to_KE] | GOBP_CELL_POPULATION_PROLIFERATION [missing_in_current_KB]<br>GOBP_MITOTIC_NUCLEAR_DIVISION [removed_from_KE]<br>GOBP_POSITIVE_REGULATION_OF_MITOTIC_NUCLEAR_DIVISION [removed_from_KE]<br>GOBP_REGULATION_OF_MITOTIC_NUCLEAR_DIVISION [removed_from_KE] |
| Event:744 Increase, Hyperplasia (Leydig cells) | 2 | 166 | 0 | GOBP_LEYDIG_CELL_PROLIFERATION [other_added] | GOBP_CELL_POPULATION_PROLIFERATION [missing_in_current_KB]<br>HP_ABNORMALITY_OF_THE_LEYDIG_CELLS [removed_from_KE] |
| Event:759 Increased, Kidney Failure | 492 | 7 | 1 | HP_ACUTE_KIDNEY_INJURY [added_to_KE]<br>2. HP_CHRONIC_KIDNEY_DISEASE [added_to_KE]<br>3. HP_RENAL_INSUFFICIENCY [added_to_KE] | GOBP_REGULATION_OF_RENAL_SYSTEM_PROCESS [removed_from_KE]<br>GOBP_RENAL_SYSTEM_PROCESS [removed_from_KE] |
| Event:459 Increased, Liver Steatosis | 4 | 1063 | 29 | (none) | GOBP_LIVER_MORPHOGENESIS [removed_from_KE]<br>HP_ABNORMALITY_OF_THE_LIVER [removed_from_KE] |
| Event:1686 Deposition of Energy | | | | (none) | GOBP_ENERGY_HOMEOSTASIS<br>WP_ENERGY_METABOLISM |
| Event:1421 Activated, LXR | | | | (none) | KEGG_PPAR_SIGNALING_PATHWAY [removed_from_KE] |
| Event:1913 Endothelial cell dysfunction | | | | (none) | GOBP_ENDOTHELIAL_CELL_ACTIVATION [removed_from_KE] |
| Event:741 Increase, Adenomas/ carcinomas (follicular cell) | | | | (none) | KEGG_PATHWAYS_IN_CANCER [removed_from_KE] |

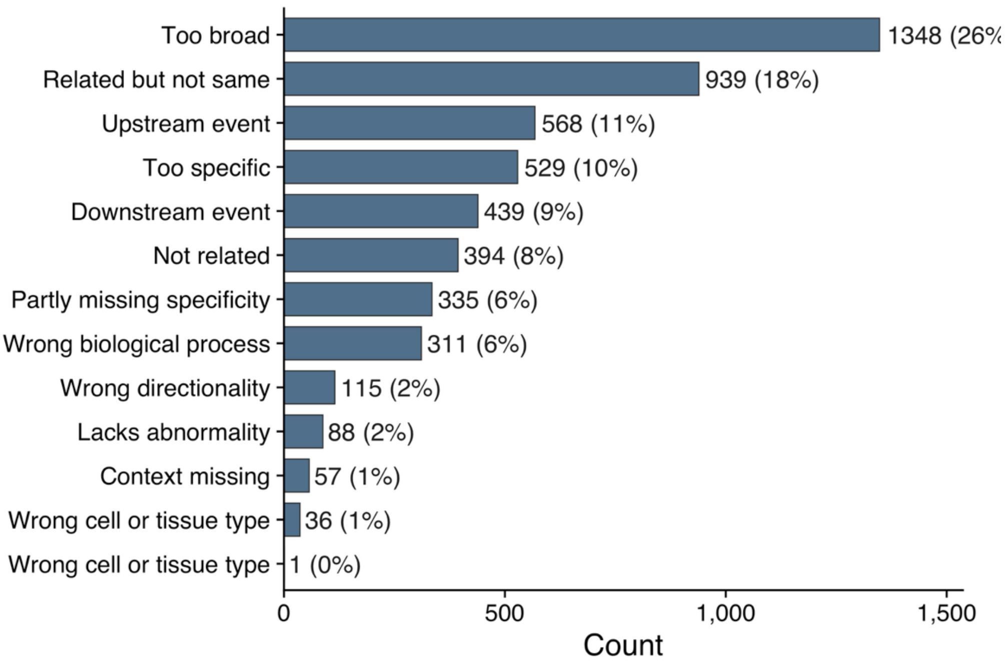


*Figure S4: Most frequent curator reason codes contributing to rejected KE-term candidates.*

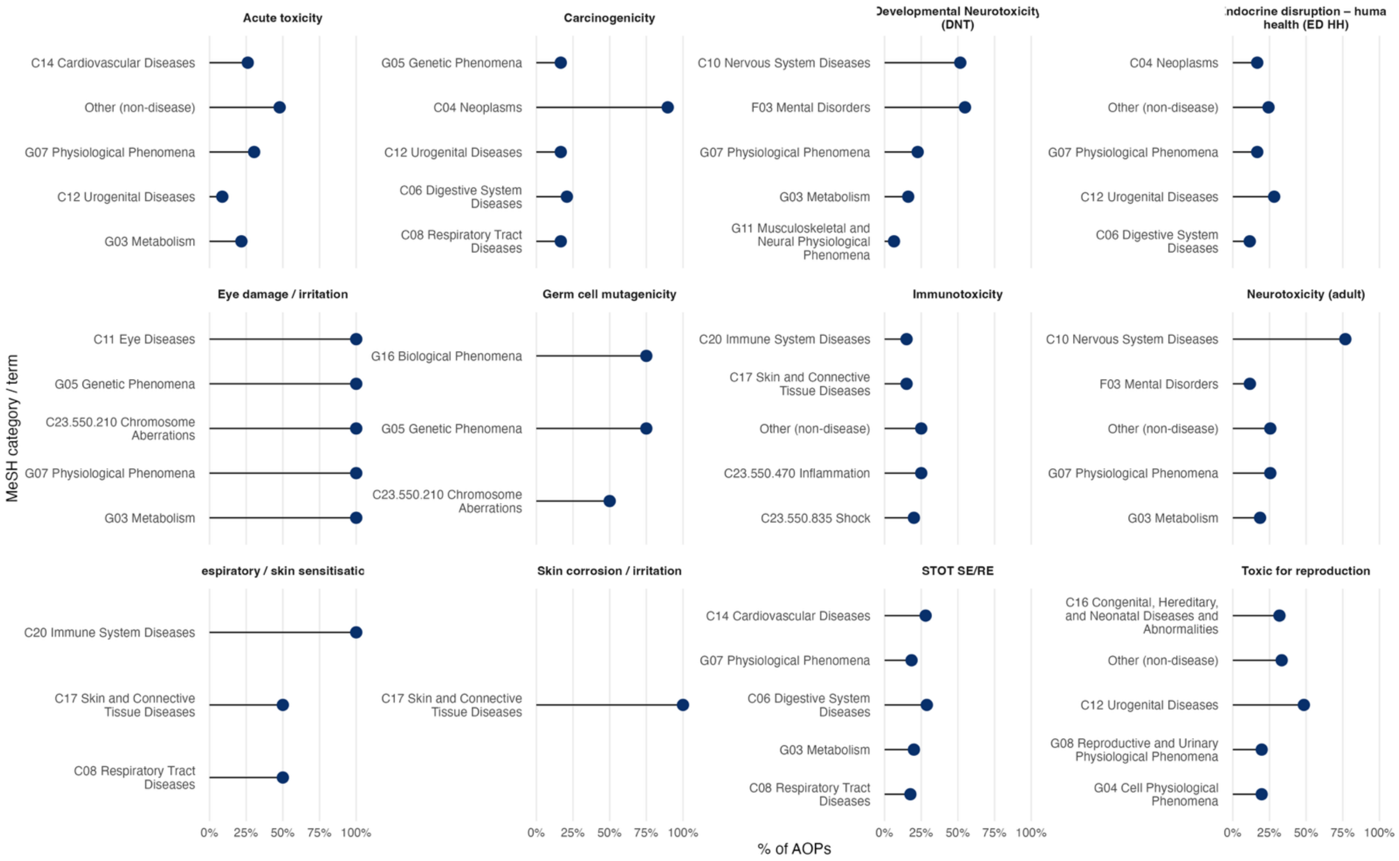


*Figure S5: Top 5 MeSH categories within each human-relevant hazard class. MeSH categories are ranked by the percentage of AOPs containing at least one mapped KE for the corresponding regulatory endpoint.*
